# Constructing future behaviour in the hippocampal formation through composition and replay

**DOI:** 10.1101/2023.04.07.536053

**Authors:** Jacob J.W. Bakermans, Joseph Warren, James C.R. Whittington, Timothy E.J. Behrens

**Affiliations:** Wellcome Centre for Integrative Neuroimaging, University of Oxford, Oxford, UK; Department of Applied Physics, Stanford University, Stanford, CA, USA; Wellcome Centre for Human Neuroimaging, University College London, London, UK; Sainsbury Wellcome Centre for Neural Circuits and Behaviour, University College London, London, UK

## Abstract

Hippocampus is critical for memory, imagination and constructive reasoning. However, recent models have suggested that its neuronal responses can be well explained by graphs, or state-spaces, that model the transitions between experiences. Here we use simulations and hippocampal recordings to reconcile these views. We show that if state-spaces are constructed compositionally from existing building blocks, or primitives, hippocampal responses can be interpreted as compositional memories, binding these primitives together. Critically, this enables agents to behave optimally in novel environments with no new learning, inferring behaviour directly from the composition. We predict a role for hippocampal replay in building and consolidating these compositional memories. Importantly, due to their compositional nature, replay can construct states the agent has never experienced - effectively building memories of the future. We test these predictions in two datasets by showing that replay events from newly discovered landmarks induce and strengthen new remote firing fields. These memories, built in replay, are compositional. When the landmark is moved, replay builds a new firing field at the same vector to the new location. Together, these findings provide a framework for reasoning about compositional memories, and demonstrate that such memories are formed in hippocampal replay.

---

A recent spate of hippocampal models suggest that hippocampus represents a state-space and its transitions, i.e. a cognitive map (Stachenfeld, Botvinick, and Gershman 2017; Piray and Daw 2021; George et al. 2021; Whittington et al. 2020). These models come in two flavours, those that infer the state-space from sequences (George et al. 2021; Whittington et al. 2020), and those that use the state-space for reinforcement learning (RL) (Piray and Daw 2021; Stachenfeld, Botvinick, and Gershman 2017). Together they explain many key hippocampal findings; from associative learning in neuroimaging studies (Garvert, Dolan, and Behrens 2017; Schapiro et al. 2016) to precise cellular responses during spatial sequences, such as place (O’Keefe 1976) and grid cells (Hafting et al. 2005). Additionally, they account for latent state representations in reinforcement learning tasks that require non-spatial behaviour such as splitter cells (Frank, Brown, and Wilson 2000; Wood et al. 2000) in spatial alternation tasks or lap cells (Sun et al. 2020) in tasks that require counting (George et al. 2021; Whittington et al. 2020). Overall, these models’ successes have been greatly suggestive that hippocampus builds state-spaces from sequences and may use these state-spaces for RL.

However, these new ideas about state-space inference seem at odds with key principles of hippocampal function that are supported by a wealth of empirical evidence. Most strikingly, hippocampus’ primary role is one of memory (Eichenbaum and Cohen 2014; Scoville and Milner 1957). This memory is part of a constructive process that also supports imagination, and scene construction and understanding (Addis, Moscovitch, and McAndrews 2007; Hassabis and Maguire 2007; Mullally, Hassabis, and Maguire 2012; Rosenbaum et al. 2009; Tulving 1985). This evidence suggests that the hippocampal representation is compositional, binding cortical information together to build a representation of the current experience. A door to the north-west. A wall to the south. A friend sitting at the table. Models of scene construction in hippocampus have already incorporated such representations ((Becker and Burgess 2000; Bicanski and Burgess 2018)). Indeed, empirically, hippocampal neurons respond to conjunctions of external features (Komorowski, Manns, and Eichenbaum 2009), as if binding together elements of a composition, and RL state-space tasks predominantly rely on hippocampus when new state-spaces are initially constructed (Packard and McGaugh 1996). The big computational benefit to compositional scene-construction and episodic memories is being able to understand and respond to situations in *one-shot*. This flexibility is missing from the traditional state-space models. For those models, learning how states relate to each other requires observing state transitions. This learning of the state-space often requires much experience, and can be brittle to policy or local transition changes (Russek et al. 2017). While these symptoms can be alleviated, for example by offline replay that performs credit assignment when new rewards or barriers are observed (Mattar and Daw 2018; Sutton and Barto 2018), the diagnosis remains the same. This raises a significant puzzle - how do the state-space models relate to the well-known memory and construction machinery of hippocampus?

Here, we unify the two through a model of *state-space composition*. To be compositional, the model of the world must be decomposable into representational sub-blocks (*z* = [*z^1^*, *z^2^*, *z^3^*, …]), where the dynamics of each sub-block are independent from one another 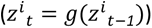. This means that new configurations of the sub-blocks (corresponding to a new world model) will have predictable dynamics with no extra learning. Instead of learning state-space transitions from experience, these transitions can now be inferred. In this scenario, the world-model for any particular situation specifies how the building blocks combine in the current world - a role we propose for hippocampal place cells ^1^. For example, composing a map of space with an object centric map (or wall- or door-centric), means the hippocampal state-space knows where the object (or wall or door) currently is in space. This feature of hippocampal compositions offers several advantages over and above hippocampus being just state-spaces or memories alone, and provides new insights into hippocampal phenomena. We show that 1) Conjunctive hippocampal cells can be reinterpreted as compositionally binding together multiple maps/variables, accounting for hippocampal single-unit responses; 2) There is a dramatic performance gain (versus standard RL) when hippocampal state-spaces are compositions of *already learned* building blocks, since policies learned on one hippocampal composition generalise to novel compositions; 3) Latent learning can be understood as building compositions in the absence of reward, so that optimal behaviour can be achieved when rewards are discovered; 4) Replay can compose states-spaces offline into memories that improve future behaviour either by updating policies or consolidating existing memories; 5) This constructive function of replay affords precise predictions of how, when, and where replay changes hippocampal representations. Indeed, when testing these predictions in single-unit recordings in rodent hippocampus, we find that 1) replay induces and strengthens hippocampal place fields: new place fields emerge after replay events, and ratemap changes align on replayed locations, 2) these ratemap changes reflect structural elements as well as rewards, and 3) the resulting responses generalise compositionally.

## RESULTS

### Model of hippocampal compositions

We propose that hippocampus makes use of reusable building blocks that it can compose together to understand new situations - just like scene construction (Fig 1a). For clarity, we explain this model in terms of spatial representations, but it applies to any situation where the world-dynamics can be split into compositional parts. In this section, we give a high-level overview of the model and its main features, which we will further elaborate on in each of the following sections. The Methods section and Supplementary Material provides further implementational details.

**Fig 1.**
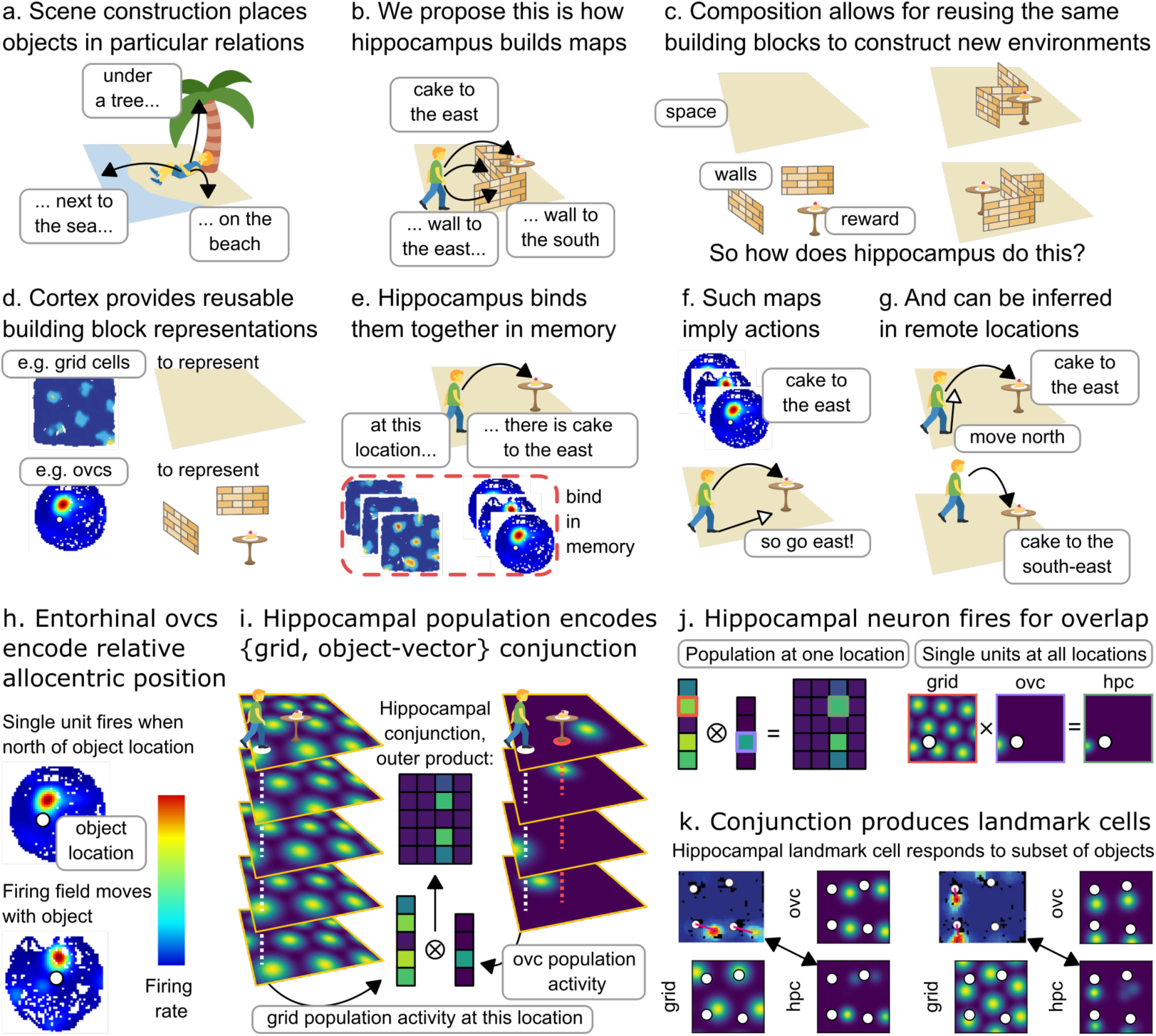
Model of hippocampal state-space composition. **a**. To construct a scene, like this (imagined) experience of being on the beach, hippocampus binds objects into relational configurations. **b**. We propose that hippocampus similarly composes state-spaces from structural elements like walls and rewards (black arrowheads denote allocentric vector relations). **c**. The advantage of composition is that new situations, like new spatial environments, can be constructed from the same building blocks. **d**. Cortex can provide such reusable representations, for example in the form of grid cells (Hafting et al. 2005) and object vector cells (OVCs; (Høydal et al. 2019)) in spatial environments. **e**. Hippocampus can construct new situations from cortical representations by binding them together in relational memory. **f**. Because these compositional state-spaces are built from reusable elements, understanding from one environment generalises to others. In particular, these maps immediately imply actions (white arrowheads). **g**. To construct the state-space, building block representations can be propagated to remote locations - online, but also offline in replay. **h**. Object vector cells (Høydal et al. 2019) encode the global relational knowledge that supports action inference. If this example cell fires, and the object is a reward, you should go south. **i**. Hippocampus then combines this information, for example from the 4 object-vector cells for reward on the right, with a spatial code, for example from the 5 grid cells on the left, into a new conjunctive memory encoded in 20 hippocampal cells, for example through an outer product. **j**. The outer product describes the population activity; on the single-neuron level, this produces place responses in hippocampus - these are place cells that carry reward information. A single unit in hippocampus (green outline) is activated when its grid cell input (red outline) and object-vector cell input (purple outline) overlap. **k**. The same mechanism can produce hippocampal landmark-vector cells (empirical: top left (Deshmukh and Knierim 2013), simulated: bottom right) that respond to some objects in the environment but not to others. These two examples show how the conjunction of the grid cell (bottom left), with firing fields on a hexagonal grid, and an object vector cell (top right), with firing fields a fixed vector relation to each object (white circle), produce a hippocampal response (bottom right) that is similar to an empirically measured response (top left).

In space, to construct a scene of a simple room, hippocampus may bind together a representation of where you are in space, *x*, with where you are relative to walls (*W*), salient objects (*o*), and rewards (*r*) (Fig 1b). This means that hippocampal cells will be a conjunction of representations for space, walls, objects, and rewards (*x*, *W*, *o*, *r*). Critically, the subcomponents (*x*, *W*, *o*, *r*) are reusable since any room is a different configuration of space, walls, objects, and rewards. Thus, the hippocampal state-space can be built immediately, for any new environment, out of different compositions of these building blocks (Fig 1c). This idea of composition through hippocampal conjunctions is not new; indeed, composing space with *observations* has been shown to support relational inference and structural generalisation (Whittington et al. 2020). But there is a crucial difference in the conjunction of space and *structural building blocks* proposed here: these structural building blocks afford generalising *policies*. The resulting hippocampal representations do not just map the world - they specify how to act in it.

A key question, however, is what should the building blocks look like? Fortunately, for space, walls, objects, and rewards (*x*, *W*, *o*, *r*), biology already tells us. In entorhinal cortex and hippocampus, along with place and grid cells that code for space, there are vector cells that point towards walls, objects, and rewards: border-vector cells, object-vector cells, reward-vector cells (Gauthier and Tank 2018; Høydal et al. 2019; Lever et al. 2009; Solstad et al. 2008). Each cell provides a distance and direction to the border/object/reward, and each population of vector cells for border/object/reward provides a map that can be path integrated (updated with respect to actions taken), just like grid cells for space, but rather with each map being centred around the border/object/reward (Fig 1d). In other words, each population of cell types represents a coordinate system, e.g. grid cells are a global 2D coordinate system, object vector cells are also a 2D coordinate system but locally centred on objects, etc. We posit non-spatial building blocks will have coordinate system representations too, i.e., vector cells but in non-spatial coordinates (Nieh et al. 2021). Lastly, these cells generalise and so are reusable - an object vector cell in one environment is also an object vector cell in another environment, with the same true for grid cells and border vector cells.

The reusability of these building blocks means any understanding from one configuration can be generalised to new configurations. This is particularly powerful for reinforcement learning, as now an agent does not have to learn a new policy from scratch for every new environment (like RL on conventional hippocampal state-spaces, e.g. SR (George et al. 2021; Stachenfeld, Botvinick, and Gershman 2017)). Instead, the compositional state-space already implies actions (Fig 1f). A reward vector cell says head towards the reward, a border vector cell says don’t crash into the border and so on. Indeed, rats (Chamizo, Rodrigo, and Mackintosh 2006) and pigeons (Blaisdell and Cook 2005) use such vector relationships to rapidly infer policies in novel environment configurations, and the temporal coding hypothesis suggests that these vectors can even be temporal rather than spatial (Arcediano and Miller 2002; Barnet, Cole, and Miller 1997). In RL terminology, credit assigned to these compositional building blocks in one situation is a useful assignment in new situations as well, i.e. the hippocampal state-space comes ‘pre-credit assigned’. This means online RL is dramatically reduced and often not necessary at all.

But how can this composition be achieved in the first place? Given the compositional state representation, the policy can be generalised - but upon entering a new environment, it must first be constructed. That requires binding representations of (*x*, *W*, *o*, *r*) at every location in the environment and in the appropriate configuration, i.e. binding the vector representation saying 3 steps north to the reward to the spatial representations when you are actually 3 steps south of the reward. This is easily achieved by binding the reward vector representation, *r*, at the current timestep to the spatial location representation, *x*, at the current timestep and storing it as a hippocampal memory {*x*, *r*} (Fig 1e). Therefore, in principle the compositional state-space can be constructed step by step in online behaviour. However, performing compositions like that is slow to propagate information throughout the whole state-space, because it requires visiting all states of the new environment. It is also error prone as it relies on path integration (Etienne and Jeffery 2004; Mittelstaedt and Mittelstaedt 1980). Fortunately, biology provides a potential resolution in the form of offline replay (Foster and Wilson 2006), which can bind building blocks together in remote locations (Fig 1g). This is both more data efficient and reduces potential errors. In this context, replay is effectively performing credit assignment since it constructs the state-space for future successful behaviour; next time we are at that remote location we already know about the reward. This function of replay also has implications for what the content of replay should be: the best replays are those that improve the map the most. We can therefore predict patterns of replay and behaviour from this principle.

### Place cells that embed global knowledge in local representations

The constructive nature of hippocampal function suggests a particular interpretation of hippocampal place cells - a conjunctive representation (Komorowski, Manns, and Eichenbaum 2009; Manns and Eichenbaum 2006), where hippocampal cells bind existing representations into a new relational configuration. At its simplest this means combining sensory input with the grid coordinate system, as suggested by previous sequence models (Whittington et al. 2020), but much more is required for it to be a useful state-space for inducing behaviour. The local (at a given location) representation must contain global (about other locations) relational knowledge (Fig 1f). Such a representation can also be built from conjunctions, but they must be conjunctions between cortical cells that encode this global relational knowledge, such as object- and border-vector cells in the medial entorhinal cortex (Fig 1h). With such a representation, unlike with existing place cell models, many transitions are inherited, rather than learnt, as they are implied by the particular combination of cortical inputs.

So what do these conjunctive representations look like? Like previous work, we assume that the hippocampal population encodes the outer product of the representations it composes together. However, previous work has formed sensory-sensory conjunctions (Kumaran and McClelland 2012), or sensory-structural conjunctions (Whittington et al. 2020), but here we form conjunctions of different structural elements (such as grids and vectors to walls or reward). This means that the state space itself is composed (Fig 1i). The resulting single-unit responses exhibit spatial tuning just like place cells (Fig 1j) - but these are place cells that carry reward information. This type of hippocampal response has been found empirically in the form of landmark vector cells (Deshmukh and Knierim 2013), which have place fields at vector relations for some objects in the environment but not others (Fig 1k).

Notably, such conjunctive representations need not be the only representations in hippocampus, and can be further tuned by learning as the environment becomes familiar, or behaviour is overlearned (for example, with an algorithm that builds a successor representation). Their power comes from their potential to generate behaviour immediately in a new environment.

### Compositional codes facilitate zero-shot behaviour

To explore this potential for zero-shot generalisation we compare two agents (Fig 2a). In both cases, we train a feedforward network *f* through supervised learning on ground-truth optimal policies to predict optimal actions *a* given a state representation *s*. In the first agent (Traditional) states are unrelated to the features across environments: they only represent space *x*. In the second agent (Compositional), states are compositions of vector cells to environment features: they combine walls, objects, and rewards (*W*, *o*, *r*). We do not include space (*x*) as part of this composition as it is not required for action selection (but we will need the spatial component later to build compositional maps in memory). In each case, we train on multiple environments with walls, objects and rewards, placed at random. The optimal policy is defined as the local actions that minimise the number of steps to reward from each location.

**Fig 2.**
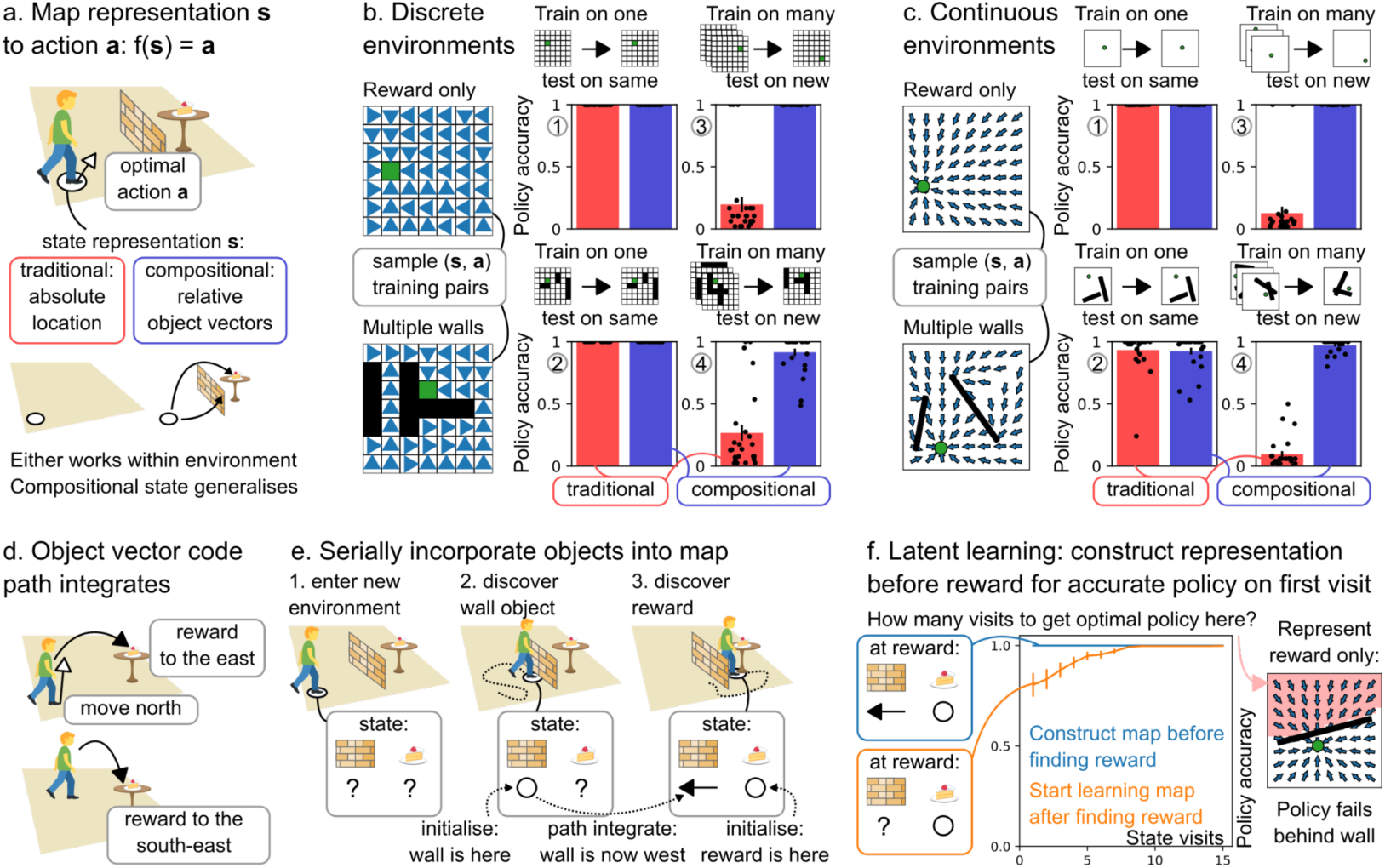
Compositional state representations generalise and can be built through latent learning. **a**. We learn a mapping *f* from state representation *s* to optimal action *a* (white arrowhead): *f*(*s*) = *a*. For a given location, *s* represents the absolute location in the environment (traditional; red; *s* = [*x*]) or the relative vector codes (black arrowheads) for all objects (walls and rewards) in the environment (compositional; blue; *s* = [*W*, *r*]). **b**. In discrete graph environments, we sample (state representation, optimal action (blue triangles)) training examples, both in simple environments with reward only and in complex environments with multiple walls. When trained on a single environment and tested on the same environment, both the traditional and compositional state representations provide accurate policies (panels 1 and 2). Only mappings from compositional state representations yield accurate policies when tested in a new environment (panels 3 and 4). Error bars: s.e.m. **c**. In continuous environments, where locations are continuous coordinates and actions are continuous directions, we find the same results. Either representation works within environments (panels 1 and 2), but only the compositional state representation generalises (panels 3 and 4). Error bars: s.e.m. **d**. How to obtain this compositional state representation in new environments? The vector code (black arrowhead) that makes up these representations path integrates. Because it follows relational rules independent of the environment, the representation can be updated with respect to the agent’s action (white arrowhead). For example, if a reward is to the east, and the agent goes north, the reward is now to the south-east. **e**. Path integration allows for serially incorporating objects (or walls or rewards) into the compositional map. The object-vector code is initialised on object discovery, and then carried along as the agent explores the environment. **f**. That means that the agent learns about the structure of the environment without being rewarded. Once they find a reward, this latent learning allows access to optimal actions on the first visit to locations behind the wall (blue). Without latent learning, the agent needs to rediscover the wall to obtain the optimal policy behind the wall (orange). Error bars: s.e.m.

In each simulation, we randomly generate a set of environments and calculate optimal policies. We then sample (state representation, optimal action) pairs (*s*, *a*) as training examples to train a feedforward network that maps state representations to optimal actions through supervised learning. We evaluate the network’s performance by measuring whether following the learned policy successfully navigates to the goal from a set of test locations (Methods 4. Policies that generalise).

While the traditional agent is able to learn arbitrarily complex policies in a single environment with the reward in a fixed location (Fig 2b1, 2b2), it immediately fails when either the reward or environmental features change (Fig 2b3, 2b4). This is unsurprising as it is not re-trained when the environment changes, and the state representation does not carry useful information across environments. By contrast, the compositional agent immediately generalises behaviour. Given the state representation contains reward vectors, this is trivially true for changes of reward location in an otherwise empty environment (Fig 2b3). However, it also holds for policies that require complicated trajectories avoiding multiple walls (Fig 2b4). These findings generalise across both discrete (Fig 2b) or continuous (Fig 2c) state and action representations. Together, these results emphasise that if hippocampal cells form a state-space for future learning, it is advantageous for them to build on existing relational structure (e.g. from cortical building blocks), as opposed to being learnt from scratch as is commonly assumed.

Compositional representations therefore allow for dramatic behavioural generalisation. In this view, the role of place cells is to bind together cortical representations into a memory, such that when the animal next visits this location they will be able to reactivate cortical representations that link to optimal actions. Paradoxically, because the agent does not need to have taken the action to build the memory, this is a memory of *future* behaviour. This memory of the future permits zero-shot inferences.

### Latent learning and laying down memories of the future

Compositional reasoning therefore changes the computation required to produce good behaviour in a new environment. Instead of learning a new behavioural policy, we must lay down new memories, but critically we must lay down these memories *everywhere* in the environment. This is important because encountering a new wall should not only affect the policy at adjacent states. Instead, the presence of a wall will change the optimal policy everywhere. When the agent is next at a remote location, its state representation must reflect the fact that there is a wall between the agent and the reward.

Like credit assignment in RL, we therefore have to update state information at remote locations. But the content of these updates is structural information rather than reward expectations. Instead of value, we must transfer the newly discovered compositional features to each state in the environment. Critically, in compositional worlds the independent parts of the representation can be updated independently. In space, object-centric representations can be path integrated independently of allocentric representations (Fig 2d). If an agent is one step east of a reward and takes a step east, it is now two steps east of the reward. Each representation is determined by the last representation and the current action. Hence one approach is simply to keep track of the path integrated representation of each object (or wall or reward) it has encountered, as it traverses the environment. However, building these representations into memories alleviates the requirement to keep track of many variables at the same time. This is possible because the representation is compositional, which means that environmental features can be added one at a time.

This process of serially integrating information into a compositional state representation naturally accounts for results in latent learning. Here animals who have experienced the structure of the environment without rewards can rapidly develop optimal policies upon discovering rewards for the first time (Blodgett 1929). Similarly, when our compositional agent explores the environment, it builds an increasingly complete state representation *s*, incorporating objects as they are encountered (Fig 2e). Then the agent finds a reward. In a simulation where the agent has already learned about the wall-vector representation, it only needs a single visit (to add the goal-vector to *s*) to any other state for access to the optimal policy (Fig 2f, blue). Without this latent learning, the agent needs to explore the whole environment again to accumulate the full state representation *s* (Fig 2f, orange).

In this simulation, we consider an agent that has explored an environment with a wall when discovering reward (Methods 5. Latent learning). To demonstrate the utility of latent learning, we compare our agent to an agent that only starts learning about the environment’s objects after discovering the reward. The former has incorporated the wall in its state representation already and will have access to the optimal policy on the first visit to a new location, including those behind the wall. The latter needs to rediscover the wall, and will not know how to optimally avoid it until then (Fig 2f, right).

### Replay builds memories efficiently

In the previous section we argued that path integration allows for serially incorporating objects into a compositional map during online exploration. However, this explanation of latent learning seems flawed. If the agent must path integrate representations by physically traversing the environment, and can only path integrate one or a few representations at a time, then it will need to traverse the environment many times to build a latent representation. To avoid this problem, rather than physically path integrating vector representations, the agent can *imagine* path integration in replay (Fig 3a). In this interpretation, replay can achieve credit assignment by exactly the same mechanism as it uses to build memories, because credit assignment is achieved by binding cortical representations into hippocampal memories.

**Fig 3.**
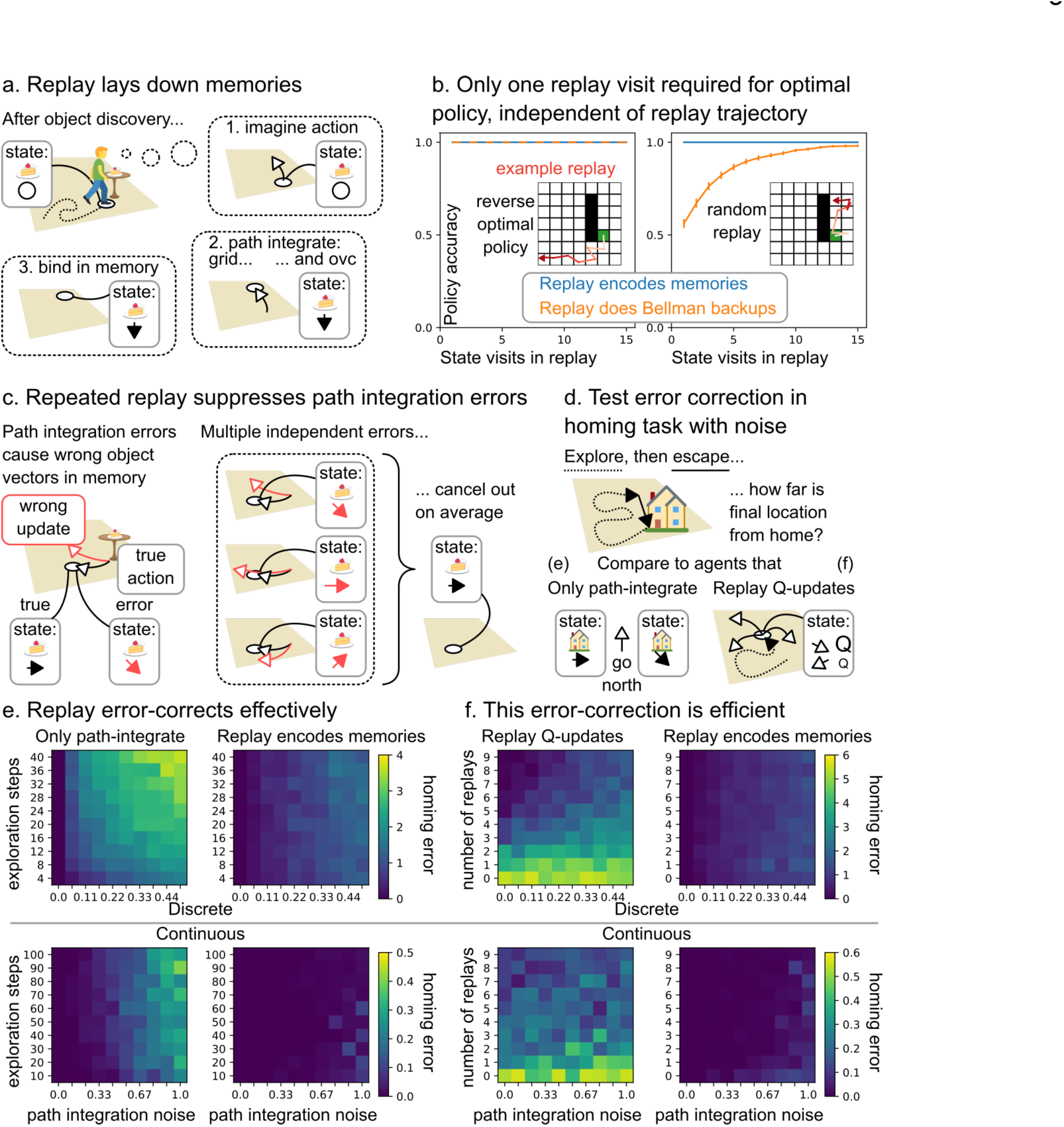
Replay builds compositional maps in memory. **a**. After discovering an object (or wall or reward), replay builds the compositional map in memory by path integrating the vector code and the grid code and binding them together to form new hippocampal memories at remote locations. **b**. Such replay (blue) provides optimal policies in a single replay visit. On the other hand, replay that performs Bellman backups for credit assignment (orange) requires a single visit for optimal actions *only* if the replayed trajectories follow the opposite optimal policy (left) but needs many more replay visits for random replay trajectories (right). Error bars: s.e.m. **c.** Path integration is noisy. This can be mitigated by forming memories in multiple different replays since path integration errors are independent and average out. **d**. We test the error-correcting capacity of replay that encodes memories in a homing task with path integration noise. The agent starts from home, explores the environment, then needs to escape back home as quickly as possible. We compare the agent that encodes memories in replay to an agent that only path integrates a homing vector (panel e) and an agent that replays Q-updates to learn which actions are expected to lead towards home (panel f). **e**. For the agent that path integrates only (left), the homing error (distance from home after escape) increases for longer exploration and higher path integration noise. For the agent that encodes home-vector representation memories in replay, the error is greatly reduced. **f**. The agent that encodes home-vector memories in replay needs fewer replays to achieve lower homing errors than the agent that replays Q-updates.

This leads to clear untested predictions. Replay events should happen in the vector cells when an animal discovers a new environmental feature (such as an object or a reward). But critically, the replay events must bind object-centred representation to their correct locations in allocentric space. This means that allocentric representations must also path integrate in replay, simultaneously with the object-vector cells. A natural candidate for this allocentric representation is the grid cell representation, which is known to path integrate (Burak and Fiete 2009; Sargolini et al. 2006). We therefore predict that replay events will involve simultaneous replay of grid cells and object-vector cells, where both cell populations replay to the same locations but in two different coordinate systems - global and object-centric respectively (Fig 3a). As the conjunction of the two will be encoded in hippocampal memory, hippocampal replay trajectories should be coherent with these (as observed in (Ólafsdóttir, Carpenter, and Barry 2016); however, (O’Neill et al. 2017) reported independent replay). This also implies the resulting hippocampal conjunctions (in this case a hippocampal object-vector-grid conjunction; landmark cells) at given locations can be active before ever physically visiting that location. Empirically, a landmark cell can be detected from its activity during online behaviour. We predict that some landmark cells will appear in replay, after discovering the corresponding object, before they appear in physical navigation. This is an example of ‘preplay’, as replay activity correlates with the response to future novel experience (Genzel et al. 2020; Dragoi and Tonegawa 2011).

Importantly, with noiseless path integration, the optimal new policy is constructed with a single visit to each state, whatever the trajectory of the replay (Methods 6. Constructive replay). This is unlike alternative interpretations of replay. For example, if replay performs Bellman back-ups (as in the dyna algorithm (Sutton 1991)) it can only update the value of any state by comparing to its neighbour, so is exquisitely sensitive to the trajectory of the replay. To achieve convergence with a single step would require prescience (Fig 3b, left) as it would need to play out (in reverse) the new optimal policy that results from the updated values. Unlike Bellman back-ups, replaying for compositional memory requires only a single visit to each state, even if the trajectory is random (Fig 3b, right). This is because path integration through a world model (spatial or otherwise!) is trajectory-independent, and so the elements that get composed at each state are identical regardless of the replay’s trajectory.

However, the reliance on path integration introduces a vulnerability: path integration is noisy, and errors accumulate over time. Replay will therefore build noisy, but unbiased, memories, where the noise can be reduced with repeated replays (Fig 3c). One attractive feature of this proposal is again that it aligns replay’s role across credit assignment and memory. Here, multiple replays are required to consolidate the existing memory (Carr, Jadhav, and Frank 2011; Karlsson and Frank 2009).

We simulate a noisy homing task where the agent explores an arena starting from a home location, until it encounters a threat and needs to return home as quickly as possible (Fig 3d; Methods 6. Constructive replay). Without replay, the homing error for the path integration agent increases both for higher noise and for longer exploration, as path integration errors accumulate (Fig 3e, left, top for discrete domains, bottom for continuous). With replay, homing errors of both causes are dramatically reduced (Fig 3e, right). Notably, Bellman backups are also susceptible to path integration errors as backups can be attributed to incorrect states. This can be ameliorated by sampling replay repeatedly (Fig 3f, left). Nevertheless, when replay is instead used to build compositional memories, smaller homing error is achieved with fewer replays (Fig 3f, right).

### Optimal replay creates memories where they matter most

Requiring as few replays as possible for accurate behaviour is beneficial as replay takes time, and time is often needed for online behaviour (Agrawal et al. 2022; Jensen, Hennequin, and Mattar 2023). Making the best use of each replay means prioritising certain replay trajectories over others (Igata, Ikegaya, and Sasaki 2021). Intuitively, the highest priority replays are those that improve behaviour the most. This intuition has been formalised in the RL framework, where replay explicitly assigns credit to states through value backups, to successfully explain a wealth of empirically observed patterns of replay (Mattar and Daw 2018). Constructive replay performs *implicit* credit assignment by binding object-centric representations, that come with pre-learned policies, to allocentric coordinates. Nevertheless, optimal constructive replay trajectories are qualitatively consistent with those predicted by the best value backups - albeit with a very different neural implementation.

Here, we define optimal constructive replay as making *those* memories that are most impactful for shortening paths towards rewards from anywhere in the environment. Such replay forms new vector representation memories, or consolidates existing ones, at locations where these representations change the policy (for example by providing a reward vector) and are frequently retrieved (for example on common paths to reward). We find optimal replays by enumerating replay trajectories and evaluating how much the resulting memories decrease the expected distance towards the goal, averaged across the environment (Methods 7. Optimal Replay).

Optimal constructive replays play out along trajectories similar to backups that maximally increase future reward, because both aim to achieve the same goal: a map that supports selecting (ultimately) rewarding actions. Updates to the map take place in locations that have the greatest impact on the overall policy, regardless of the representation that underlies that policy. Upon reward discovery, such optimal updates distribute the reward information back through the environment (Fig 4a), whether through value backups or constructive replays. Once complete, the reward map can be further improved by consolidating the existing memories: additional constructive replays suppress path integration errors, along similar trajectories as ‘need’-driven value backups (Fig 4b)(Mattar and Daw 2018). These examples illustrate that the principle of replaying whatever improves the policy most can be applied irrespective of the replay mechanism: trajectories compatible with optimal value backups are often compatible with optimal state-space construction too.

**Fig 4.**
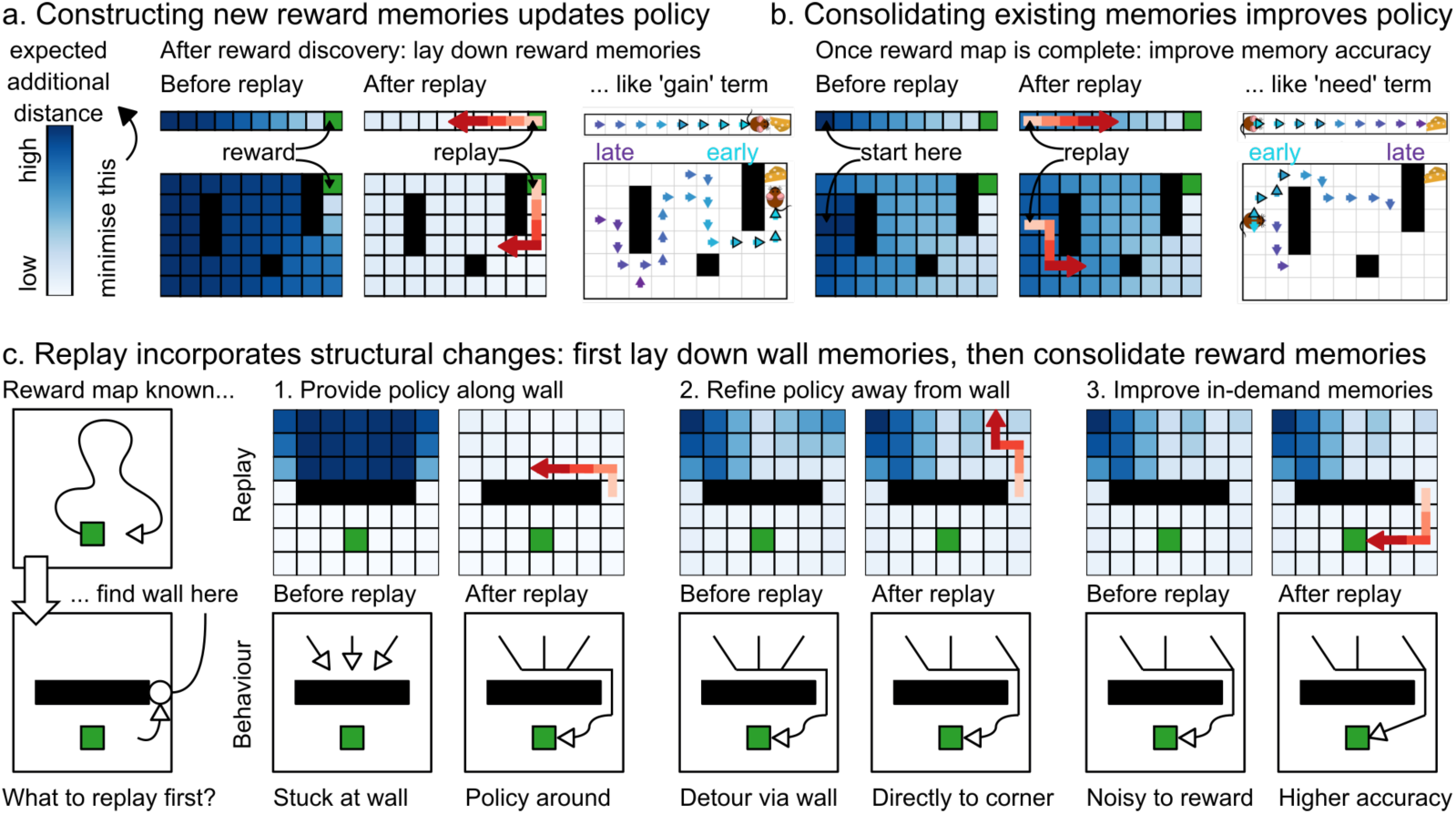
Optimal replay constructs then consolidates maps. **a.** Optimal constructive replay minimises the total expected additional distance (the sum of additional distance per state plotted in blue). Upon finding a reward, that means laying down reward memories, producing simulated trajectories similar to ‘gain’-dominated value backups (Mattar and Daw 2018). **b**. Once the reward map is sufficiently complete, optimal replays consolidate existing memories that will be needed often in the future, by encoding additional memories to error-correct path integration noise. Such simulated replays play out along similar trajectories as ‘need’-dominated value backups (Mattar and Daw 2018). **c**. Constructive replay naturally accounts for structural changes, such as the discovery of a wall in the environment. Optimal replay encodes memories where they matter most. In this situation, that produces simulated replays first along the wall to guide the policy around the wall, then away from the wall for a more direct policy, and then towards reward to improve the accuracy of memories that will be retrieved many times.

However, that replay mechanism does sharply distinguish the two. Value backups treat the map, whether in hippocampus or cortex, as static, and change striatal synapses to reflect state values. Constructive replays change the hippocampal map itself. A new (replayed) composition that combines a particular vector and grid code recruits a new hippocampal representation - the conjunction between the two. That means that, in addition to reward changes, constructive replay also naturally accounts for structural changes (Fig 4c). We simulate replays when an agent discovers a new wall in a familiar environment with a rewarded location. Optimal replay first runs along the wall (opposite side to the reward) while encoding wall-vector memories. Now all paths behind the wall are successfully rerouted around the wall, though they still go directly to the wall first, and then around. To make these trajectories less bendy, and more efficient, the next replay updates state-representations progressively further away from the wall, so that paths end up going directly to the edge of the wall. Finally, optimal replay consolidates the reward memories that are retrieved most often: those on the path from wall to reward. While no recordings of replay events exist in such situations, exactly these progressively less bendy behavioural trajectories are observed in behaviour when barriers are placed between a start and goal (home) location in a homing task (Shamash et al. 2021).

### Empirical evidence of replay for map making

But a more precise empirical validation of our model requires testing our (preprint-preregistered (Bakermans et al. 2023)) neural predictions. In particular, we’ll test whether “some landmark cells will appear in replay (…) before they appear in physical navigation” in two datasets (Methods 8. Testing replay predictions). We find that upon replay, changes in the ratemap of hippocampal neurons 1) closely align to the neuron’s replayed location, 2) reflect structural changes beyond just reward, and 3) generalise in a compositional way. Together, these results support a role for replay in building maps (1) from structural elements (2) through composition (3). We demonstrate how replay changes maps in hippocampal recordings during an alternating home-away well task (Pfeiffer and Foster 2013). We find many replays while the animal sits at the home well, where planning is not very useful (the away well is in a random location) but mapping how to return home later is. Indeed, when we decode these replay trajectories, we discover that ratemap changes occur exactly where a cell fires in replay. Moreover, for certain cells these ratemap changes generalise across home well locations, affording compositional reconfiguration when the environment changes. Finally, such environment changes may include structure beyond reward: in a four-room maze task where doors lock halfway through the experiment (É. Duvelle et al. 2021), we find new place fields that emerge after replay at a recently locked door.

We probe how replay changes maps in hippocampal recordings collected by Pfeiffer & Foster (Pfeiffer and Foster 2013) in an alternating home-away well task (Fig 5a). This task involves an arena where reward appears in hidden reward wells, following a very specific pattern: first at a fixed ‘home’ well, followed by a random ‘away’ well, then at the same fixed home well again, then a different random away well, et cetera. The animal thus needs to alternate between random exploration to discover the currently rewarding away well, followed by memory-guided navigation to return to the home well. According to our model, this task can be solved through a hippocampal home-well map that provides this return policy. Our theory proposes that this map is constructed from reusable vector representations in replay. We therefore need to test how replay changes hippocampal representations. This paradigm is ideal for that: the animal returns to the home well many times, providing ample opportunity for map-making replays; meanwhile, the search for the random away well yields good behavioural coverage of the arena, to probe what that map looks like.

**Fig 5.**
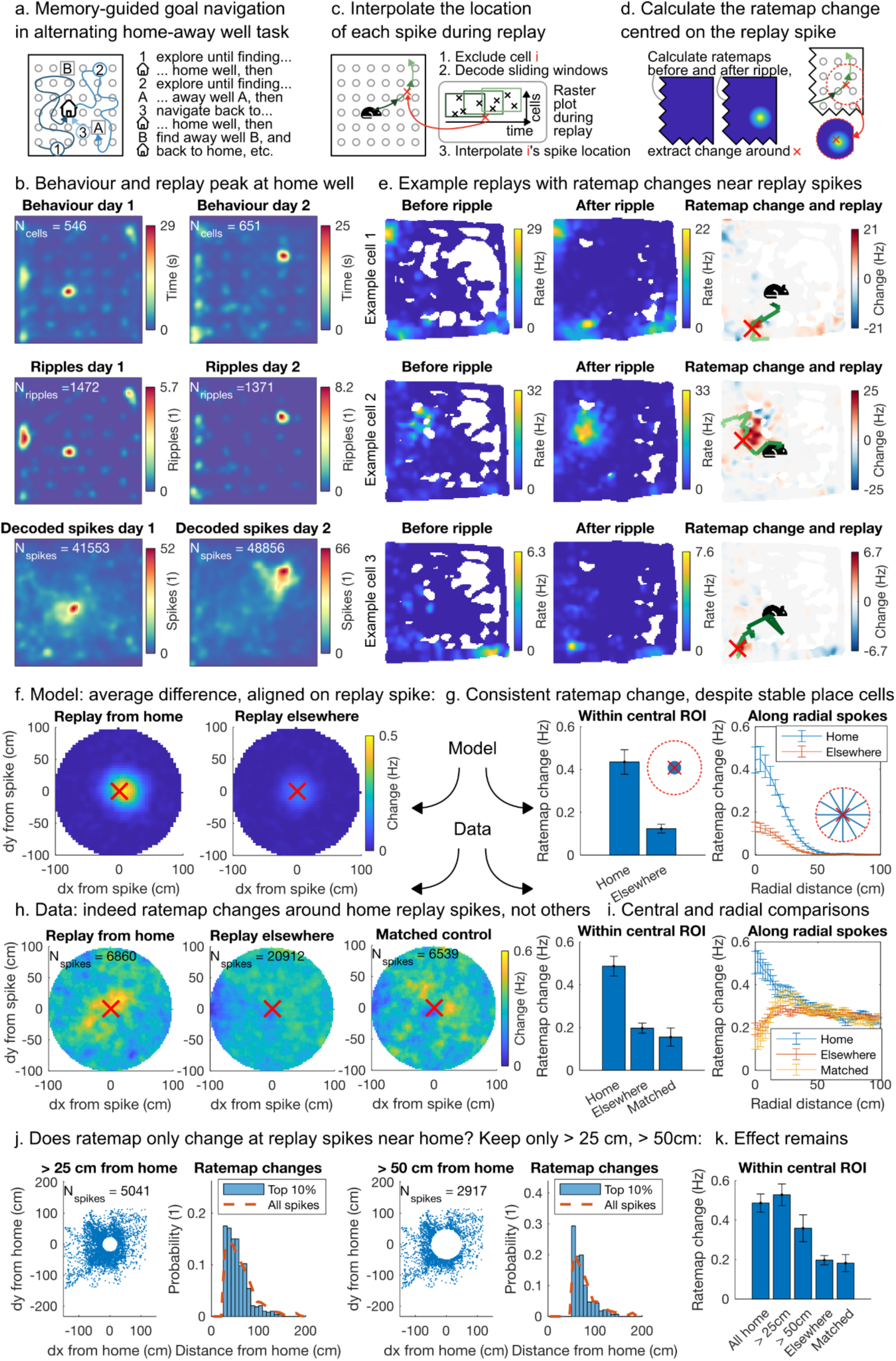
Replay changes hippocampal rate maps. **a.** In the alternating home-away well tasks, the animal needs to alternate between exploring for the currently rewarded random away well and navigating to the remembered stable home well. **b**. The occupancy averaged across animal peaks at the home well location (first row), and many sharp wave ripples occur there (second row). Within the ripples, interpolated spike locations are distributed across the arena (third row). These spike locations are calculated as follows: **c.** We estimate the location of a spike of cell i during replay by decoding the replay trajectory while excluding cell i, then interpolating the spike time along the replay trajectory. **d.** We then measure the ratemap before and after that replay event for a cell, and calculate the ratemap change aligned on its replay spike location. **e**. Three example cells with ratemap change (right column; red: positive, blue: negative change) near the interpolated replay spike (red x) along the decoded trajectory based on all other cells (green line, from dark to light). **f.** In a population of simulated hippocampal neurons, we average ratemap changes aligned on replay spike locations across neurons and replays. We find a consistent change at the replay spike that is stronger for replays from home that create new landmark fields (left) than replays elsewhere that do not (right). **g.** Averaging the aligned simulated change maps within a 8cm region of interest at the replay spike location (left), or along radial spokes extending out 100 cm in equally sampled direction from the replay spike location (right), further illustrate the same effect. We find a consistent ratemap change despite the inclusion of place cells with stable ratemaps. Error bars: s.e.m. **h**. In the same analysis on empirical data rather than simulations, we find the same result. On average the ratemap changes align on the replay spike for home replays (left), but not for replays elsewhere (middle), also after matching them for time within the experiment (right). **i.** Averaging these aligned change maps within a narrow central region of interest (left) or along radial spokes (right) again highlight this effect. Error bars: s.e.m. **j.** It is not just the replay spikes near home that contribute to these ratemap changes, as evidenced by excluding replay spikes within 25cm (left) or 50 cm (right) from home. Some of the largest ratemap changes occur in remote locations. **k**. After keeping only remote (>25cm, >50cm) replay spikes, the effect of ratemap changes at the replay spike remains.

We identify replay events by detecting sharp wave ripples in the local field potential. We first observe that, although previous analyses of this data have focused on ripples during navigation (Pfeiffer and Foster 2013), in fact many ripples occur when the animal is sitting at the home well (Fig 5b). Are these ripples modifying the hippocampal map in a compositional way? We are going to examine whether neural firing fields during open field behaviour change after a replay, and where that change happens. If replay is indeed responsible for creating new conjunctions, we expect a neuron’s ratemap to change at the exact location where that neuron fired in the replay event because that the replay event is hypothesised to path integrate relational knowledge around the map. To find out, we not only need to detect replay events, but also decode their contents. More specifically, we need to know where a cell fired in a replay event, in a way that is not contaminated by its own ratemap. To probe what happens during such events, we decode the animal’s position from their neural activity in sliding windows through the replay. But we do this decoding in a slightly convoluted way: we repeat the decoding for each neuron that spikes during the replay event, while excluding that neuron, then interpolate the location of its spike (Fig 5b). This gives an estimate of where a neuron spikes in replay that is independent of its own rate map. So how does the ratemap change - what is the difference in the neuron’s ratemap before and after the replay event - around that location (Fig 5c)?

We find a strong effect of aligned positive ratemap changes at the replay spike location, specific to replays from the home well. There are many of these as the animal frequently replays while it sits at the home well. When we apply the procedure described above to such replay events, we discover hippocampal cells that form a new place field right around the interpolated spike location (Fig 5e; green line: decoded replay trajectory, red cross: interpolated spike location). To understand how such emerging place fields look across a population of recorded neurons, we simulate replay-induced map changes in a synthetic neural population of hippocampal place cells (Fig S9c) and landmark cells (Fig S9a,b) (Methods 9. Home-well mapping simulations). Our simulation shows that for replays that build maps, on average we expect a small but consistent increase in firing rate right at the location of the replay spike (Fig 5f, left). The increase is consistent because of our simulated landmark cells, which first fire in replay and then obtain a new receptive field. But the increase is small because it is averaged across many neurons (including our simulated place cells, that do not change their ratemaps) and replays (whereas only the first replay creates a new field from scratch). Nevertheless, the resulting aligned change will be greater for replays from home that lay down vector representations, than for replays elsewhere that do not (Fig 5f, right; Fig S10). We show the same effect in a different way by averaging the changes within a 8 cm radius around the replay spike (Fig 5g, left), and by averaging across spokes in each direction from the replay spike (Fig 5g, right).

Indeed, when we average the ratemap change around the replay spike across cells and replay events in Pfeiffer and Foster’s dataset, we find a strong ratemap increase right at the replay spike – for replays from home, but not for other replay events (Fig 5h; matched control: selection of replay events elsewhere so that their time distribution over the session matches that of the replays from home). There are two reasons to consider these replays elsewhere as control events. The first is that they may be used for planning, rather than memory modification, as in the original interpretation of this data. The second is that they may consolidate memories through error correction of noisy initial maps, as suggested in modelling sections above. In this case, replays elsewhere will still induce some changes to the landmark cells, but these will be much smaller than the changes made when the fields are laid down in replays from home. Averaging the resulting change maps within a narrow region around the replay spike, or along radial spokes outwards from the replay spike, further illustrates these differences (Fig 5i). This effect holds even when only considering replay spikes more than 50cm from the home well (Fig 5j,k; Fig S7).

But these responses could have been just reward-related, because there is reward at the wells. Our theory posits that the hippocampal map composes other elements too - in fact, the previous section predicted wall-representation replays upon wall discovery. We therefore test if replay also maps such structural elements – here: blockades – in recordings by Duvelle et al. (E. Duvelle et al. 2020) in a four-room configurable maze task (É. Duvelle et al. 2021). In this task, the animal needs to navigate to one of four rooms connected by doors, then forage there, and then navigate to a different room, et cetera. Halfway through the experiment, some doors lock, without changing their appearance: the only way to find out if a door is locked, is by trying it. A locked door has major implications for the maze topology, so we expect the animal to incorporate this change in the hippocampal map through replay on discovery.

We collect ratemaps before and after the doors close, and look for replay when the animal discovers this structural change (Fig S6a). Because we can’t detect SWRs or decode replays in this dataset, we define replay at doors through ‘non-local door spikes’ (Fig S6b). A non-local door spike occurs when a neuron fires outside any of its place fields while the animal sits at a closed door, and serves as a proxy for a replay that propagates the obstacle-vector. We identify cells in which such replay precedes the emergence of new place fields (Fig S6c), and find that across the population, cells that replay are more likely to get new place fields (Fig S6d; linear regression, one-tailed t-test; Fig S7). Because of the indirect replay measure here, this second set of results should be interpreted with some care; they are suggestive but to be conclusive will require a dataset where enough cells are recorded simultaneously to decode replay trajectories.

The previous results support a role for replay in map-making. For this map to be compositional, cells need to generalise their home response across environments (Fig 6a). For example, in the dataset from Pfeiffer & Foster a home-vector cell would fire south-east of the home well, regardless of its location (Pfeiffer & Foster change the home location each day). Hippocampus can then recombine such representations to construct new maps of any home-well configuration by binding these vector representations to specific locations. This conjunction produces hippocampal landmark cells (Fig 1k) with firing fields at the same spatial relation for the home well on day 1 and day 2. Can we find evidence of these generalised home responses?

**Fig 6.**
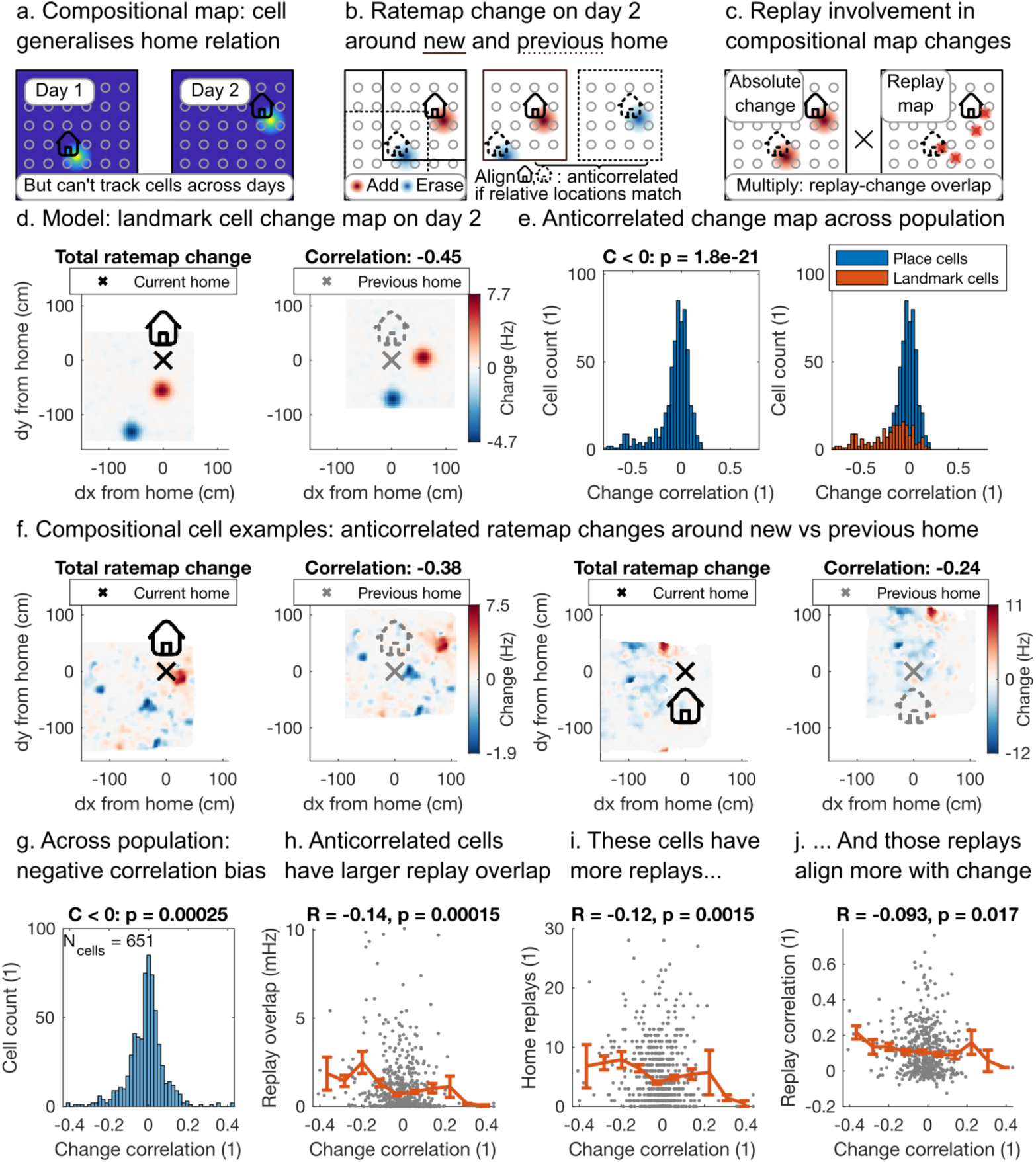
Ratemap changes generalise. **a.** In a compositional map, home responses generalise, so that new home-well configurations can be mapped by recombining existing elements. **b**. To detect generalised responses without access to cells across multiple days, we plot the ratemap change of a cell on the second day, and ask if the change with respect to the new home is anticorrelated to the change with respect to the previous home. **c**. If replay is involved in such changes, the cell’s replay spike locations should overlap with the ratemap changes. **d.** Example ratemap change on day 2 of a simulated landmark cell whose object-vector cell fires to the south of the home well. When we align the same change map first on the new home well location (left, black) and then on the old home well location (right, grey), the positive and negative changes overlap so the maps anticorrelate. **e.** Across the population of simulated hippocampal neurons, that produces a correlation histogram with a negative bias (left). When we split the contributions of simulated place and landmark cells (right), we find that the landmark cells are the source of this negative bias. **f.** Two examples of recorded neurons with anticorrelated changes (red: positive; blue: negative) with respect to the new (left, black) and previous (right, grey) home. **g.** Across the population, this ratemap change correlation is negatively biassed, suggesting a subpopulation of neurons with generalised home responses (one-tailed t-test against zero). **h.** The ratemap changes of those anticorrelated neurons overlap more with replay spikes (linear regression, one-tailed t-test; grey: cells; orange: average of y-values within 10 bins along the x-axis): **i**. they have more replays, and **j**. their change locations correlate more with their replay locations. Error bars (g-j): s.e.m.

Not only do we find neurons that show such responses, we also discover that for the same neurons, replay correlates with ratemap changes. We cannot track cells across days, but we can probe the cumulative change of each neuron’s ratemap on day 2 (Fig 6b). If a neuron generalises, it will create a new place field at a fixed spatial relation to the new home – but it should also erase the place field at that same relation to the previous home. That produces a measurable signature: the ratemap change aligned on the current home should anticorrelate with the ratemap change aligned on the previous home. Our simulation of neural responses to the home well on different days recovers this signature of generalised home well responses. The example simulated landmark cell in Fig 6d fires when the animal is to the south of the home well. Therefore the increase in firing rate in red on day 2 (Fig 6d, left) aligns with the decrease in firing rate in blue on day 2 with respect to where the home well used to be on day 1 (Fig 6d, right), resulting in a negative correlation of the shifted change map. Our simulation shows that this produces a negative bias in the correlation coefficients (Fig 6e, left), due to the consistent remapping of landmark cells (Fig 6e, right).

Indeed, we find neurons with anticorrelated change maps (Fig 6f), and on the population level there is a negative bias in the aligned change correlation (one-tailed t-test; Fig 6e). To investigate replay involvement, we also ask whether the location of replay spikes for a neuron overlap with these ratemap changes (Fig 6c). It turns out that these cells on the left of the distribution are also the ones with the highest replay overlap (linear regression, one-tailed t-test; Fig 6f) – they have both more replay spikes (Fig 6g) and the locations of replays and ratemap changes correlate (Fig 6h) – which suggests that replay may be responsible for these changes. Taken together, these results indicate that within the population of hippocampal cells with spatial responses, there is a subpopulation of landmark cells (the ones that receive input from entorhinal vector cells, according to our model) that exhibits vector responses that may be constructed in replay.

### Composing non-spatial and hierarchical building blocks

Constructive replay thus carries structural information beyond just reward to compose representations that imply actions in remote locations. Importantly, this works equally well for non-spatial building blocks. That is important because empirically, hippocampus supports non-spatial reasoning (Aronov, Nevers, and Tank 2017; Dusek and Eichenbaum 1997), for example through representation of hierarchy (Kjelstrup et al. 2008; Shapiro, Tanila, and Eichenbaum 1997) and context (McKenzie et al. 2014). Computationally, non-spatial composition opens up a whole range of new problems to which the same principles can be applied. These problems need to a) decompose into building blocks so a common function *f*([*z^1^*, *z^2^*, *z^3^*, …]) = *a* maps states to optimal actions, and b) allow independent forward models for each building block 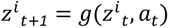 so building blocks can be replayed separately and then recombined in memory. Path integration is a specific spatial example of such forward models; more generally, they can be learned to capture a broad variety of environment dynamics.

We now show that our model works in a hierarchical task that mixes space and non-space (Methods 10. Non-spatial replay). This task consists of multiple spatial rooms (one of which contains a reward) joined together into a (non-spatial) loop by doors, but where the doors are at random spatial locations within each room and behave like teleports (i.e. the locations of the doors are unrelated to the rooms they transition to; Fig 7a). We provide a state representation that composes both spatial within-room vector codes, towards doors (*d^1^*, *d^2^*) or reward (*r*), and a non-spatial between-room vector code, that specifies the number of rooms between the current and the rewarding room (*c*). Importantly the *d^1^*, *d^2^* and *r* representations are reused across different rooms. As before we train a neural network *f*([*d^1^*, *d^2^*, *r*, *c*]) to predict optimal actions on tasks with many different room sizes and door locations, and test on an entirely new hierarchical environment. We see that the network generalises to unseen configurations and provides accurate policies in each room towards the rewarded room (Fig 7b). This highlights generalisation across the hierarchy as anything learned in one room applies in another; alternative approaches like hierarchical RL would need to learn separate policies (or *options*) in each room.

**Fig 7.**
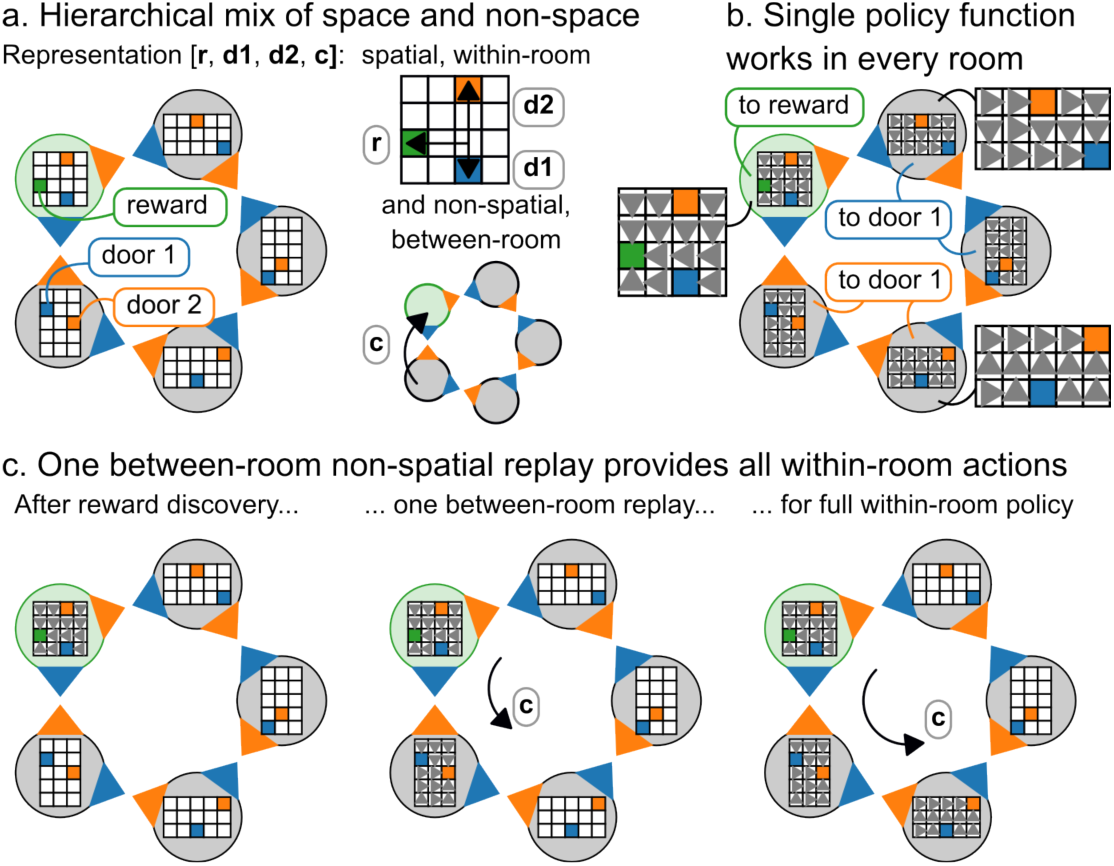
Policy and replay over hierarchical non-spatial building blocks. **a.** An example hierarchical environment that mixes space and non-space, consisting of rooms on a non-spatial loop. One room contains reward (green); each room contains two doors (blue and orange) at random locations that transition the agent to the next room along the non-spatial loop. We provide a full state representation that consists of within-room spatial vectors to reward and doors (*r*, *d^1^*, *d^2^*) and a between-room non-spatial vector code (*c*) to the rewarding room. **b**. A policy mapping *f*([*d^1^*, *d^2^*, *r*, *c*]) = *a* learned across many hierarchical environments provides optimal actions in unseen new configurations, using the same within-room vector codes (*d^1^*, *d^2^*, *r*) in each room. **c**. Upon finding a reward, and replaying its within-room location, a single between-room replay propagates the rewarding room-vector code, *c*, thus providing the optimal policy to each of the other rooms.

In the situation described above, we provided the state representation. An agent, however, can only initialise a reward vector code (or rewarding room code) after actually observing a reward in one of the rooms. Similarly to before, we use replay to propagate this knowledge within the rewarded room (i.e. replay *r* vectors), but additionally replay across rooms (via *c*). This replay is thus decoupled over space (*r*) and non-space (*c* vectors). Since the replay of *c* is at a hierarchical level, a single replay suffices to obtain the optimal policy in all un-rewarded rooms (Fig 7c). The non-spatial between-room vector *c* can be viewed as a contextual signal that modulates behaviour depending on the state of the environment (like chopping or frying, depending on the current stage of a recipe). And because its forward model can implement arbitrary non-spatial dynamics (such as the non-spatial loop), it could guide actions in game environments that are currently challenging for AI agents (Wang et al. 2021).

## DISCUSSION

Recent results have suggested a path towards building a formal understanding of neural responses in flexible behaviours. By assuming that the hippocampal formation builds a state-space (cognitive map) from sequential observations, these models have not only revealed computational insights into the hippocampal involvement in reinforcement learning (Piray and Daw 2021; Stachenfeld, Botvinick, and Gershman 2017), but also accounted for a variety of single neuron responses (George et al. 2021; Whittington et al. 2020). However, while this has the potential to bring formal explanations to an array of new scenarios it is not clear how these models relate to classic hippocampal functions such as episodic memory (Addis, Moscovitch, and McAndrews 2007; Scoville and Milner 1957), scene construction (Hassabis and Maguire 2007), or imagination. Indeed, hippocampal patients are impaired not only in navigation, but also in scene recognition (Graham, Barense, and Lee 2010) and imagination of future and fictitious scenes (Hassabis et al. 2007; Mullally, Hassabis, and Maguire 2012; Rosenbaum et al. 2009). In this work, we propose a framework where these disparate functions can be expressed in the same formal language. We formalise hippocampal state-spaces as compositions of reusable building blocks. We have shown that this affords flexibly generalising behaviour to new situations on first encounter, by simply rearranging previously learned components.

We propose that hippocampal cells provide conjunctions of pre-learnt building blocks, specifying their arrangement in the current experience. This allows us to reinterpret several hippocampal phenomena and offer new computational roles for others. For example, hippocampal place and landmark cells are conjunctions of grid cells with sensory cells (Whittington et al. 2020) and object-vector cells respectively. This means that hippocampus no longer needs to learn transitions itself, as these are inherited from the building block dynamics. While scene construction may be relevant for representing individual states too, what we propose here goes beyond that: it is about constructing the whole state space. Composing a state space affords inferring transitions, rather than learning them from sequential observations. Furthermore, we show that forming memories of these compositions during explorations, provides the ideal state-space for *future* behaviour when observing the reward (latent learning). Importantly, if the building blocks have a forward model then memories can be formed offline in replay. This enables an agent to efficiently build a compositional state-space for future behaviour. The idea that replay serves future behaviour has already proven influential (Mattar and Daw 2018). Here we show how a different underlying mechanism can provide rapid zero-shot credit assignment, and hence generalisation. This constructive interpretation of replay naturally extends to changes in task structure as well as reward, and integrates these ideas with the compositional nature of hippocampal/entorhinal representations.

Our empirical data support this compositional mechanism in two complementary datasets. In Duvelle et al. (É. Duvelle et al. 2021) the data are many examples of first discovery of new landmarks (here closed doors), allowing us to show that cells with new firing fields were likely to have fired in replay when the door was discovered. In Pfeiffer and Foster (Pfeiffer and Foster 2013) there were fewer first-discovery events but many visits to the landmark (here a home-well). Furthermore, due to high cell count we were able to decode exactly where replays visited. We showed that each replay event changed the place map at exactly the inferred location where the cell fired during replay. This was only true for replays at the home well and was true for fields many tens of centimetres from the home well. Furthermore, we showed that these changes are compositional (with the caveat that pure vector responses in hippocampus would produce the same signatures; entorhinal recordings would be needed to prove conjunctive coding). When the landmark moves, replay builds a new place field at the same relative location, but now to the new landmark. Together these findings suggest that when a new structural element is discovered in the world, replay lays down a compositional memory that embeds the vector to this landmark. We note that an earlier version of this paper on biorxiv (Bakermans et al. 2023) included this as a pre-registered test of the theory. We would like to thank the reviewers for encouraging us to perform this test.

Computational models, in particular REMERGE (Kumaran and McClelland 2012) and TEM (Whittington et al. 2020), have previously shown that conjunctive coding in hippocampus supports generalisation. However, the type of conjunction and the scope of generalisation as a result are fundamentally different here. REMERGE learns links between sensory features within an environment through sensory-sensory hippocampal conjunctions. That allows for sensory generalisation, like transitive inference, within the environment, but never across environments, because there is no abstract state-space structure embedded in the hippocampal representation. TEM does learn abstract state-space structure to make sensory-structural hippocampal conjunctions, which allow for sensory generalisation across environments. But in our model the state-space structure itself is compositional. By forming structural-structural conjunctions, we compose state spaces from structural building blocks. As a result we can generalise behaviour: we make inferences about actions, not sensory input. Rather than zero-shot predicting what the agent will see, as in TEM, we zero-shot predict what the agent will do. Because that requires knowledge of global relational structure (e.g. a wall to the east, and a reward to the south), this global information must be propagated to the local hippocampal representation. That is exactly what we propose replay is for.

Importantly, we do not propose that all composition in the brain requires hippocampus. Language, for example, is compositional but is not impaired after hippocampal damage; the ability to construct complex motor skills from motor primitives similarly remains intact. In fact, even generalising policies from structural building blocks here does not require hippocampus: the building blocks are represented in cortex. But hippocampus becomes essential when these cortical representations need to be stored in relational memories. This is exactly the same machinery as episodic memory or scene construction, or other compositional tasks that hippocampus is involved in: hippocampus constructs new representations of whatever the relevant inputs are. In the case of combining structural building blocks here, the resulting representations form compositional maps. Critically, this allows us to take advantage of another hippocampal capacity: replay. Hippocampal replay can propagate memories of combinations of structural building blocks around the map, which support optimal future behaviour. In summary, not all composition takes place in hippocampus, but the hippocampal machinery for relational memory and replay turns out to be remarkably powerful for making compositional maps that generalise policies.

It is notable that in our model the state-space is simply the combination of appropriate *cortical* building blocks. The role of the hippocampus in the model is one of memory and replay. Access to this memory/replay system prevents the model from having to continually track all the elements of the forward model. Instead it can track one at a time (adding to existing state representations stored in memory) and can do so in replay rather than during behaviour (making online behaviour instantaneous). This minimal role for hippocampus is appropriate when worlds can be composed perfectly from existing primitive building blocks. By contrast, in models where hippocampus learns the transition structure (George et al. 2021; Stachenfeld, Botvinick, and Gershman 2017), any transitions can be modelled (after learning). One intriguing possibility is that both systems are at play. Compositional inference gives rapid flexible approximate behaviour, which is nuanced by modelling of transitions. In such a system another possible role for replay is to build new cortical primitives when experiences have been poorly modelled by the existing repertoire.

While we have elucidated a role for replay in the online setting, this framework also suggests a neural interpretation of the two previously proposed roles for replay in sleep (Ellis et al. 2020): building cortical primitives (as above), and learning the policy on the compositional state-space. Here replay could help learn compositional policies by generating training examples for policy mapping *f*. By sampling random vector representations for walls and rewards, the agent dreams up arbitrary new environment configurations. Instead of encoding memories, it then replays to simulate trajectories in those imagined configurations. A successful rollout that reaches reward provides a training pair of the sampled vector code with an optimal action. This is a natural extension to the idea of learning an inverse model during sleep (Helmholtz machine (Dayan et al. 1995)), but it is particularly powerful for compositional forward models as “sleep training” can include samples that have never been experienced (Ellis et al. 2020).

Notably, we do not learn any building block representations in this paper, but instead assume representations in the form of a grid code for space and vector codes for the other building blocks. The ideas in this paper are not limited to grid and vector representations but apply to any forward models that can be decomposed into sub-models where actions have independent consequences. It will be particularly powerful to learn these forward models from experience - another possible role for sleep replay. Critically, our model assumes that these representations can be bound into hippocampal conjunctions such that any state in one elemental model can be bound with any state in another (the wall can be at any (*x*, *y*)-location). For this binding to work, it is likely that different elemental models should be expressed in different neural populations (disentanglement). Indeed, previous work has also shown that this is also the most energy efficient form of representation (Whittington et al. 2022).

Our idea of constructing behaviour from reusable building blocks is related to meta-RL. In its broadest sense, meta-RL proposes a system that slowly learns over many episodes to train a fast within-episode learning system (learning to learn (Harlow 1949)). Our compositional building blocks, and function *f*, permit fast within-episode learning, and we have assumed the building blocks are learned over many episodes. The implementation of meta-RL that has so far captured neuroscience, however, is where the dopamine system slowly trains prefrontal recurrent networks to solve sequential tasks (Wang et al. 2018). The key difference between our proposition is that we utilise conjunctive hippocampal representations (and Hebbian memories) for *explicit* building block compositions, whereas presumably a form of implicit composition is taking place in the meta-RL prefrontal models. Lastly, we note that transformer neural networks (Vaswani et al. 2017), the state-of-the-art meta-learners (Adaptive Agent Team et al. 2023), are computationally closely related to hippocampal compositions (Whittington, Warren, and Behrens 2022).

Interestingly, while we have not explicitly tried to solve hierarchical RL, our framework nevertheless provides ingredients that may prove important. Hierarchy splits a big problem into smaller subproblems which may share structure and require similar solutions; this is exactly what our policy function *f* learns from building block codes, since they naturally generalise across subproblems. This is unlike the options framework (Botvinick 2012) and related approaches like (Saxe, Earle, and Rosman 2017), which only partially exploit hierarchical structure, as each smaller subproblem still needs to be solved individually. Additionally, because representations in our model are factorised between levels, new pieces of information about one hierarchical level immediately permeate to all other levels.

Extreme generalisation by composition is a fundamental property of human and animal cognition. Whilst this has been self-evident in cognitive science for many decades, it is typically ignored in computational neuroscience. In this work, we have taken this notion seriously and tried to align it with a series of properties of hippocampal function; some long-known - memory, construction and conjunctive coding; and some more recently discovered - state-spaces for controlling behaviour. As the community attempts to find formal descriptions for computations underlying increasingly rich and complex behaviours, we believe that compositional reasoning will play an increasingly important role in future models, not only of cognition but also of neural responses.

## Acknowledgements

We thank Brad Pfeiffer, David Foster, Éléonore Duvelle, Roddy Grieves, and Hugo Spiers for sharing their datasets and for their assistance in navigating those. We very much appreciate that Duvelle et al. have made their well-organised and carefully documented dataset publicly available (E. Duvelle et al. 2020). We thank Alon Baram and Kris Jensen for helpful comments on earlier drafts of the manuscript. We thank the following funding sources: Sir Henry Wellcome Post-doctoral Fellowship (222817/Z/21/Z) to J.C.R.W.; and Wellcome Principal Research Fellowship (219525/Z/19/Z), Wellcome Collaborator award (214314/Z/18/Z), and JS McDonnell Foundation award (JSMF220020372) to T.E.J.B.. The Wellcome Centre for Integrative Neuroimaging and Wellcome Centre for Human Neuroimaging are each supported by core funding from the Wellcome Trust (203139/Z/16/Z, 203147/Z/16/Z).

## Data availability

All new data in this study (Figure 1, 2, 3, 4, 7) were obtained through simulation, for which the code is made available at https://github.com/jbakermans/state-space-composition. Additionally, this study re-analysed existing data, collected for publication in Pfeiffer & Foster (Nature 2013) and Duvelle et al. (Current Biology 2021). The Pfeiffer & Foster dataset (Figure 5, 6) was provided on request by the authors. The Duvelle et al. dataset (Figure S6, S7) is publicly available as an online resource under doi 10.25493/7NJQ-ANH at https://search.kg.ebrains.eu/instances/Dataset/6ba32f59-e7a0-4cc3-a465-86e1fdf2ffc9.

## Code availability

Code for models, simulations and analyses is available at https://github.com/jbakermans/state-space-composition.

## METHODS

A Python implementation of all models and simulations is available at https://github.com/jbakermans/state-space-composition. Here, we first describe the modelled worlds and agents in general, and then provide details of each figure’s simulations.

### 1. Discrete and continuous worlds

We implement agents in continuous and discrete environments. Although similar in principle, these two types of agents and environments are slightly different in their practical implementation. The discrete agent behaves on a graph, whereas the continuous agent behaves in a two-dimensional Euclidean space. Both discrete and continuous settings are deterministic Markov Decision Processes (MDP).

The **discrete** environments are defined by a set of locations and actions as in a deterministic MDP. All results here assume rectangular square grid worlds, with actions ‘North’, ‘East’, ‘South’, ‘West’. We generate environments by adding walls and rewards on top of these regular grids. We represent rewards with a population of object vector cells, where each cell fires at a specific vector relation to the reward, i.e. a cell that fires 1 step to the East and so on. We represent *each* wall with two populations of object vector cells - each vector population centred on one of the wall ends. In more details, we calculate the object vector population activity at location *x* relative to object *o* by concatenating the vectors of one-hot encoded distance from *x* to *o* along each action, with -1 distance for actions in the opposite direction. For example, for an object 1 step east and 3 steps south from *x*, the representation is *concatenate*(*onehot*(−*1*), *onehot*(*1*), *onehot*(*3*), *onehot*(−*1*)). Thus at any location, there are 4 vector cells active in the whole population - one for each action. In square grid worlds this representation has redundancy, because east is the opposite of west and north the opposite of south, but this setup allows for accommodating any type of non-grid graph too.

In **continuous** environments, locations become continuous (*x*, *y*)-coordinates, and actions are steps in a continuous direction. We place walls and rewards within a square 1m x 1m arena, at random locations and orientations. Again, we represent rewards by a single population of object vector cells and walls by two populations, one centred on each wall end. A single object vector cell is defined by a 2-dimensional gaussian firing field, tuned to a specific distance and direction from its reference object; the firing fields of the full object vector cell population are distributed on a square grid centred on the object. We thus calculate the object vector population activity at location *x* relative to object *o* by evaluating the population of gaussians centred on *o* at *x*.

### 2. Agent: tracking representations

Our agent does not have access to these vector representations when it enters a new environment (except Fig 3 where we provide the full state representation). It needs to discover objects first. During exploration the agent observes its location and initialises a vector representation on object discovery, i.e. sets *concatenate*(*onehot*(*0*), *onehot*(*0*), *onehot*(*0*), *onehot*(*0*)) at that location *x*. From then on, it updates the vector representation on each transition. This update combines two components: 1) it path integrates its previous vector representation with respect to its action, and 2) it retrieves any existing vector representations previously stored in memory based upon the current observed location (the memory links vector representations to location representations). It then stores the updated representation at the new location in memory.

To implement this process we use two practical abstractions from a full neural system. First, we use a direct location signal (an id of the location), instead of a neural grid code from which location is traditionally thought to be decoded from. Second, we instantiate memory as a key-value dictionary, rather than a hippocampal attractor network that stores conjunctions (these are in fact directly relatable to each other (Ramsauer et al. 2021)). In this dictionary, the space-representation serves as the key, and the object/wall/reward-vector as value, so that stored vector-representations can be retrieved at a given location. These abstractions do not change any model principles but keep the implementation simple.

The **discrete** agent initialises an object’s vector representation when it is at a location adjacent to it. Then on each step, it path integrates the representation - but due to path integration noise, the *represented* vector relation might diverge from the *true* vector relation. We model this noise as a probability distribution *p_PI_* over the updated representation, so that it reflects either the correct transition with probability (*1* – *e_PI_*) or one of the neighbours with probability *e_PI_*. In addition to path integration, as stated above, the agent relies on memory to update its vector representation. After observing its new location, it retrieves vector representations inferred there previously. That produces another probability distribution *p_m_* over represented vector relations, with the probability of a vector representation proportional to the number of retrieved memories of that representation. The agent then samples the final updated representation from the weighted sum of the path integrated and memory-retrieved distributions: 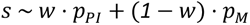. Finally, it stores the sampled representation in memory at the new location.

The **continuous** agent tracks representations in a similar fashion. It discovers an object when it comes in range (5 cm), initialises the corresponding vector representation, and from then on updates it after every step. We model path integration errors in continuous space by adding gaussian noise to the direction and step size of the update to get a path integrated vector representation *s*_%&_. Again, the agent combines this path integrated representation with a representation that is retrieved from memory. Continuous memories store vector representations at continuous locations; the agent retrieves all memories created within a cutoff distance (5 cm) from its new location, then weights the vector representations stored in these memories with a softmax over the cosine similarities between the memory location and the new location to obtain *s*_’_. The updated representation is the weighted sum of path integration and memory-retrieval: 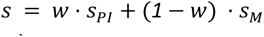. The agent then encodes this inferred representation in memory at the new location.

### 3. Conjunctive memories

We propose that hippocampal representations are conjunctions of building block representations, and that these conjunctions can be stored as memories in hippocampal weights (Fig 1). While it is possible to implement a conjunctive representation in different ways, here we use an outer product representation, i.e. the conjunction *c* between representations *a* and *b* will have cells corresponding to the product of all pairs of *a* and *b* cells. So if *a* has 3 cells, and *b* has 4 cells, then *c* will have 12 cells. To demonstrate what conjunctive representations (e.g. between spatial representations and sensory representations) would look like in hippocampal recordings, we simulate populations of grid cells, object vector cells, and sensory neurons. Each of these neurons is defined by one or multiple spatial gaussian firing fields, arranged on a triangular grid (grid cells), at fixed distance and direction from an object (object vector cells), or at a random environment location (sensory neurons). See example conjunctive representations in Fig 1.

To show how hippocampal landmark cells could result from conjunctions of object vector cells and grid cells (Fig 1k), we generate object vector cell ratemaps and perform the outer product with a grid cell representation. This leads to landmark cell responses since a grid cell’s peaks do not always align every object (or object vector cell), and so each conjunctive cell will not necessarily be active around every object.

The Tolman-Eichenbaum Machine (TEM (Whittington et al. 2020)) proposed a specific biological implementation of this conjunctive memory. The authors model place cells as conjunctions of grid cells in medial entorhinal cortex and sensory codes in lateral entorhinal cortex. They use a Hopfield network with fast Hebbian plasticity to implement the hippocampal memory. The result is an auto-associative attractor network. The recurrent dynamics of such a network allow for pattern completion of partial representations, so that one part of the conjunction can retrieve the other part. That means that observing the current state of the environment supports inference of the corresponding grid code. Or, the other way around, access to the grid code of a location allows for predicting the corresponding observation. We hypothesise a similar neural implementation of conjunctive memories here, except that the conjunctions combine structural building blocks. However, our actual simulations will mostly abstract away this biologically plausible setup. Instead, we implement conjunctive memories through a simple ‘key-value’ dictionary, where one part of the conjunction acts as the ‘key’ to index the other part of the conjunction as the ‘value’ for encoding and retrieval (also see Methods 2; Supplementary Material 1.4).

### 4. Policies that generalise

If an agent has access to the full compositional vector representation, it should be able to behave optimally even if it has never seen the particular composition before (Fig 2a-c). We demonstrate this by learning a policy mapping *f*(*s*) = *a* that maps an input state representation to an output optimal action. The compositional state representation that serves as input to this policy mapping concatenates the vector representations as described in Methods 1 for all objects in the environment. When there are multiple objects in the environment, like a reward and two walls, the representations of the objects are concatenated in consistent order, e.g. first the reward and then the two walls. That is important because a reward has different consequences for behaviour from a wall; between the walls, the order does not matter. Effectively this means the agent encodes vector relations for different objects in different populations of object-vector cells, and knows the identity of each object. An alternative but equally viable implementation could use a separate object-vector population for each object type, with multiple activations for multiple objects of the same type. For the environment with only a reward (Fig 2b1,3 and Fig 2c1,3) the input representation consists of just a reward vector code, and for the environment with multiple walls the input representation concatenates a reward vector code and three (in discrete environments, Fig 2b2,4) or two (in continuous environments, Fig 2c2,4) wall vector codes. Here, we always provide complete representations (that contain all objects in the environment) as input to the policy mapping, but if objects are missing (e.g. when they don’t exist in the current environment, or when they have not been identified yet) their representations can be set to zeros to mimic a population of object-vector cells without activity. The output action is a vector of probabilities across the four discrete actions in discrete environments; in continuous environments, the output action is a two-dimensional vector that contains the sine and cosine of the optimal direction. For the compositional vector representation, s is the concatenation of object vector population activities described above (e.g. rewards, walls etc). As a control, we also learn a mapping for a representation of absolute location (‘traditional’), i.e. a representation that does not know about walls or rewards.

We implement the function *f* as a feedforward neural network, with three hidden layers (dimensions 1000, 750, 500 in discrete environments; 3000, 2000, 1000 in continuous environments) with rectified linear activations. For the wall representation (which is two populations of vector cells per wall), we include an additional single network layer (common to all walls) that takes in the two populations and embeds them (same embedding dimension as input dimension) before feeding them in to the first hidden layer of of *f* - this is not necessary for learning, but does speed it up. We then sample [state representation, optimal action] pairs as training examples from environments with just a reward and environments with a reward and multiple walls, and train the network weights in a supervised manner through backpropagation. To evaluate a learned mapping, we sample locations and then simulate a rollout that follows the learned policy, and calculate the fraction of locations from where following that policy leads to reward (within 5cm).

To test whether policies learning from a *single* environment generalise (Fig 2b1,2; Fig 2c1,2), we sample training examples from one environment (discrete: 1000 samples; continuous: 2500 samples) and test on the same environment, for 25 environments independently. To test whether policies learned from *many* environment generalise (Fig 2b3,4; Fig 2c3,4), we train the network on 25 environments in parallel (batched input), sampling training examples in each environment (discrete: 200 samples; continuous: 500 samples) before sampling a new set of 25 environments (100 times). We then test the learned mapping for a new set of 25 environments not included in training.

### 5. Latent learning

Upon entering a new environment, we allow the agent to obtain and update compositional representations of objects/walls even in the absence of reward - this is latent learning (Fig 2e-f). To show the utility of latent learning (Fig 2e), we simulate agents with and without latent learning in a discrete environment with a wall and reward. We consider the situation where both agents have already explored the environment and found the wall, and now they have just discovered the reward. The agent that does latent learning then has a full vector representation of wall and reward, while the agent without latent learning only knows about the reward vector (as it did not obtain or update the wall representations when it saw the wall earlier). We simulate both agents as they continue their exploration. The latent learning agent can use path integration to continue updating wall/reward vector representations, and can use these to calculate the optimal policy, wherever it goes even if it has never been there before. The non-latent learning agent, on the other hand, needs to rediscover the wall to incorporate it in its state representation before it can behave optimally from all locations. To calculate the difference in their optimality, for each location behind the wall (where the full vector representation is required for appropriate behaviour), we calculate whether the agent would be able to successfully navigate to the reward based on its current representation on every encounter. We average policy success across these locations on first, second, et cetera until fifteenth, encounter and repeat the simulation 25 times in different environments.

### 6. Constructive replay

Replay offers a way of carrying vector representations to remote locations, without having to physically navigate there (Fig 3). During replay, the agent imagines actions, path integrates location (grid cell) and vector (object vector cell) representations, and binds them together by encoding a new memory of the resulting combination. We model these replayed transitions like the ones in physical navigation, with two important differences: 1) the agent cannot observe the transitioned location like in behaviour, so there can be path integration errors in the memory location (the ‘key’ in the dictionary) as well as in the representation (the ‘value’), and 2) it only relies on path integration to update representations during replay, without the memory retrieval.

First, we compare an agent that encodes memories in replay like this to an agent that instead carries out credit assignment by temporal difference learning through Q-updates (Fig 3b). We apply ‘backwards’ Q-updates (temporal discounting factor γ = *0*.*7*, learning rate *α* = *0*.*8*), in the opposite direction of replay (i.e. after a replay transition from *a* to *b* we calculate a backup from *b* to *a*), to make credit assignment more efficient. We sample replay trajectories that start from the reward location and either extend out along a random policy (Fig 3b, right), or a reverse-optimal policy (Fig 5b, left). We then calculate for each step in the replay trajectories whether the currently learned policy, either according to the encoded vector representation or the Q-values, provides an optimal path to reward from that step’s location. We aggregate policy success by the first, second, et cetera until fifteenth, replay visit to each location and average across locations, and repeat the simulation 25 times in different environments.

Then, we investigate the consequences of path integration noise (Fig 3e,f). We simulate a homing task, where the agent starts from home and initialises its home-vector representation, then explores the arena, until it needs to escape to home as quickly as possible. During exploration, the agent builds home-vector memories by replaying from the home location 5 times every 4 steps, and retrieves these memories when it is updating its current home-vector representation (i.e. using memories to reduce path integration noise). During escape, it selects actions that lead home according to its current home-vector representation. In two different experiments, we compare the agent to two controls. The first control agent only path integrates its homing vector, without encoding or retrieving any memories (and without any replay). In this experiment (Fig 3e) we vary path integration noise and the total number of steps during exploration. The second control agent carries out Q-updates to learn actions (discretised direction in the continuous environment) expected to lead home in on-policy replays, and uses the learned Q-values to find the way home during escape. Because the current location can be observed from the environment, the read-out of Q-values during escape does not suffer from path integration errors, but path integration noise does affect the replayed Q-updates: during replay, the location to update Q-values for needs to be path integrated, so credit can get assigned to the wrong place. In this experiment (Fig 3f), we vary the path integration noise and the number of replays that the agent engages in every 4 steps.

### 7. Optimal replay

Given the proposed constructive function of replay, what should its content be (Fig 4)? We define a replay as optimal if it maximally reduces the expected additional distance to reward (as compared to before replay), summed across all locations. We search over all possible replays to find the replay trajectory that best improves this metric. Conceptually, there are two ways replay can improve the metric: Either by adding vector representations of previously missing elements into the compositional map, or by consolidating already existing representations to reduce path integration noise. We now describe how we calculate the expected additional distance, and show how replay can improve it by these two ways.

To fully elucidate how the memories formed in constructive replay improve expected additional distance, we need to think about the possible representations at each location. In particular, each location either has access to the *full* state representation, composed of vector representations for all objects/walls/rewards, or a *partial* state representation, where an element of the composition is missing - but replay can provide the missing element. Partial representations can have different consequences for policy optimality. Sometimes the resulting policy is still successful (for example, when the missing element is irrelevant for the policy, like a wall to the north when there is reward to the south), but other times the agent will get stuck. Replay must then choose which locations to visit to make *partial* state representations *full* or to further improve representations that are currently noisy.

There are multiple possible ways to formalise expected additional distance. We formalise it by comparing the optimal path versus the path taken by an agent that operates in three regimes related to whether it has access to *full* or *partial* representations. The first regime is when it has a *full* representation - in which case it can path integrate straight to the goal. The second regime is when it has a *partial* representation - then it can path integrate but may get stuck in states. The third regime is random behaviour - it can switch to this behaviour after it gets stuck. This means the agent will always eventually find the reward. The agent can start using a *partial* regime then transition into the *full* regime on visiting a state with a memory of the *full* state representation, or it can transition from *partial* to random if the agent gets stuck. We formally model the above using three absorbing Markov chains on the discrete environment. The dynamics of these Markov chain are specified by three transition matrices, defined by policies as 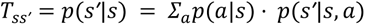: the transitions *T^full^* for the policy given the full state representation, *T^part^* for the policy given a partial state representation, and *T^rand^* for the random exploration policy. The reward is an absorbing state for each of the chains. For *T^part^* there are two additional types of absorbing state: locations where the partial policy gets stuck, for example when there is an unexpected wall in front of the reward, and locations where memories of the full state representation have been encoded. Together, these three chains allow us to analyse a process where an agent that does not have access to the full state representation follows the partial policy until it arrives at reward, arrives at a memory, or gets stuck. If it arrives at reward, it is done (1). If it arrives at a full representation memory, it switches to the dynamics of the full policy *T^full^* that takes it to reward (2). If it gets stuck, it starts random exploration following *T^rand^* until it either arrives at reward, or hits a memory that provides it with the full representation, so it can follow *T^full^* to reward (3). The total expected distance for a location to reward thus becomes the sum of the expected distances of all these scenarios, weighted by their probabilities:

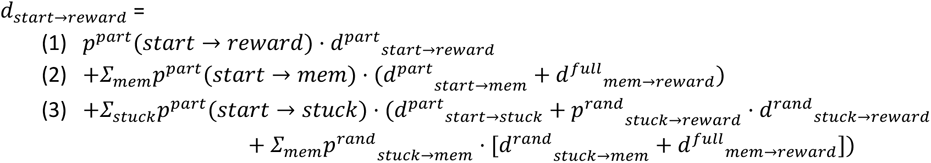

To evaluate this expected distance we need to calculate the absorption probability and expected time until absorption for a given chain and a given absorbing state. To get these, standard Markov chain analysis separates the transition matrix *T* into the dynamics between transient (non-absorbing) states *Q* and transitions from transient to absorbing states *R*, and defines fundamental matrix *N = (I_t_−Q)*^−1^, where *I*_*t*_ is the identity matrix with dimension number-of-transient-states. The probability of getting absorbed at state *j*, starting from state *i*, is given by the (*i*, *j*)-element of *B* = *NR*. The expected time until absorption *anywhere* starting from state *i* is given by the sum of the *i*-th row of *N*. To get the expected time until absorption *at a specific absorbing state k*, we calculate the sum across columns of the fundamental matrix *M* of an adjusted transition matrix *U* that expresses transition probabilities *given eventual absorption at k*: *U_ij_ =B_ik_T_ij_/B_jk_* (Clyde 2022).

The above considers replay that provides missing elements to the compositional map in memory. But because memories are noisy, due to path integration errors, replay can also improve the policy by encoding memories for elements where these already exist. Repeated replay of existing memories improves the accuracy of those memories. To model this, we additionally inject noise into the transition matrices *T^full^* and *T^part^*. Because this noise is caused by path integration errors, we model it by mixing the policy at one location with the policy of the neighbours, plus a fixed amount of random policy: 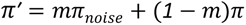. Here *πʹ* is a vector of action probabilities, *π* is the noise-free policy vector, and *π_noise_* is the mean of the policies across all neighbours and a random policy with equal probability for each action. The amount of mixing *m*, which determines the policy noise level, is lowered from 0.2 to 0.1 when replay creates an additional memory of an existing representation at that location (i.e. goes from 1 memory to 2 memories).

To actually calculate the optimal replay trajectories, we first calculate the expected distance to reward for each location, subtract the true distance to reward, then sum across all locations. Then we sample all possible 5-step replay sequences starting from the agent’s current location and recalculate this total expected additional distance given the memories created in that replay. The optimal replay is the one that decreases the total expected additional distance the most.

### 8. Testing replay predictions

We test our replay predictions - in particular: replay builds maps, or more prosaically: “some landmark cells will appear in replay (…) before they appear in physical navigation” - in two datasets: the alternating home-away well task collected by Pfeiffer & Foster (Pfeiffer and Foster 2013) and the reconfigurable four-room maze by Duvelle et al. (É. Duvelle et al. 2021).

In the Pfeiffer & Foster dataset, to define replay events, we use sharp wave ripples detected in the local field potential as provided by the authors, in combination with the population firing rates. We calculate the population firing rate as a histogram with 1ms bins of all spikes when the animal’s speed is below 5 cm/s, then smooth the histogram through a Gaussian kernel with 10ms standard deviation. A replay event starts when the population rate exceeds its (non-zero) mean before the ripple, and ends when the population firing rate drops below its mean after the ripple. We then repeatedly decode the replay trajectory during the replay event, excluding each neuron that spiked during the event once. We follow the memoryless Bayesian decoding algorithm from Pfeiffer & Foster:

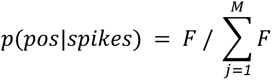

where

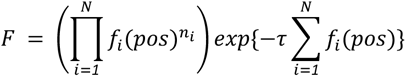

where *pos* is the animal’s position, *spikes* the neural spiking with *n*_!_the number of spikes of neuron i, *τ* the decoding time window, *N* the number of neurons, *M* the number of position bins, and *f*_i_(*x*) is the position tuning curve (or the ratemap) for neuron i. We calculate this ratemap by discretizing the arena in 2cm x 2cm bins, selecting only times when the animal moves faster than 5 cm/s, then counting the amount of time the animal spends in each bin and the number of spikes of neuron i in each bin. We smooth both resulting maps with a Gaussian kernel with a standard deviation of 4 cm, then divide the latter by the former to obtain a ratemap. We verify our decoding accuracy by calculating ratemaps from the first half of a session, then decoding position in 250 ms windows throughout the second half of the session, and comparing the decoded position to the true position (only including windows when the animal moves faster than 5cm/s) for all animals for both sessions (Supplementary Material Fig S6). To decode replay trajectories, we apply the same algorithm within a 20 ms window that we slide through the replay event by 5 ms steps. To interpolate the location of a spike of the excluded neuron, we find where the spike time falls between decoding window centres, and interpolate the spike’s position between those windows’ decoded positions accordingly. We calculate the ratemap for the neuron before the replay event and after the replay event, then subtract before from after to get the ratemap change. We translate this change map to place the interpolated spike’s location at the origin. The final product is a replay spike centred change map for a particular spike of a particular neuron for a particular replay event.

We compare these replay spike centred change maps between replays from home and replays elsewhere. A home replay meets three criteria: the animal is within 10 cm of the home well at the time of the replay, the decoded replay position is within 30 cm from the home well at least once, and the mean decoded position is at least 40 cm away from the animal’s location. These criteria make sure the replay trajectory passes by but also extends out from the home well, where the animal is physically located. A replay elsewhere meets three different criteria: the animal is more than 10 cm away from the home well, the decoded trajectory stays at least 30 cm away from the home well at any time, and the mean decoded position is at least 40 cm away from the animal’s location. These replays elsewhere function as a control condition for any results we find for the replays from home, because we do not expect replays elsewhere to change the hippocampal map like the replays from home. But there is a possibility that the replays elsewhere generally occur at different times in the experiment (for example, most home replay may occur early and most replays elsewhere late). In that case the control does not work: any results will be confounded by time. Therefore we sample a selection of the replays elsewhere that matches the home replays for when the replay event occurs in the experiment. To do so, we calculate a 5-bin histogram of the event time of the home replays, sample an equal number of replay events from the replay elsewhere and bin them the same way, and calculate the correlation between the two 5-dimensional vectors of histogram counts. If the histogram counts of the home replays and the sub-sample of replays elsewhere are correlated, that means that the distribution of replay times is similar between the two. We therefore generate 10000 samples of replays elsewhere and keep the sample with the highest correlation. Given these three groups of replay events (home, elsewhere, and matched), we calculate how the replays change ratemaps. To do that, we average the replay spike aligned change (calculated as explained in the previous paragraph) for all cells that participate in the events - if a cell has multiple spikes in one replay event, we average the change map for that cell first (Fig 5f). Additionally, we calculate the average change within an 8 cm radius for each cell in each replay of the three replay groups (Fig 5g, left) and the average change in 50 steps of 2 cm along 20 uniformly distributed angles (Fig 5g, right).

In the same dataset, we look for change maps that are anticorrelated with respect to the current home well versus the home well on the day before, because anticorrelation would mean that the cell generalises its vector relationship to the home well across home well locations (Fig 6). We average the change maps across all replay events for each cell, then translate this changemap so the current home well ends up in the origin, and translate a copy of the changemap so the previous home well is at the origin, then correlate the two. We calculate the distribution of these correlations (Fig 6e), and plot them on the x-axis (Fig 6f-h) against replay measurements on the y-axis. The first of these replay measurements (Fig 6f) is the replay-changemap overlap. If the interpolated locations of replay spikes of a neuron overlap with the location of its ratemap changes, that suggests that replay plays a role in making the changes happen. We quantify this replay-changemap overlap by calculating the absolute of the total changemap, and multiplying it elementwise by a map of interpolated replay spike locations, smoothed with a Gaussian kernel of 4 cm standard deviation, then averaging the resulting map. A high overlap can be caused both by a high number of replay spikes, or by precise alignment of replay and change. To calculate the contribution of the former, we simply count the number of home replays that the cell participated in (Fig 6g). To calculate the contribution of the latter, we calculate the replay-changemap overlap again, but this time normalise the map of replay spike locations so that it sums to 1, to isolate the spike location from the number of spikes (Fig 6h).

In the Duvelle et al. dataset, we cannot decode replay or detect sharp wave ripples in the local field potential. The number of simultaneously recorded neurons is too small for accurate decoding, and the quality of the local field potential signal is not sufficient for reliable sharp wave ripple detection. Instead, we define replays through ‘non-local door spikes’. We look for these non-local door spikes in the third of the five sessions that make up a full experiment: two sessions with all doors open, two sessions with certain doors closed, and another session with all doors open. The third session is the one in which the animal first discovers which doors have closed. We define a non-local door spike as a spike that meets three criteria: 1) the animal must be within 10 cm of a closed door, 2) moving slower than 5 cm/s, 3) and the cell cannot have a place field at the animal’s current location. We use the place fields that the authors detected in the original study, from both the second session (doors open, before animal discovers closed doors) and the fourth session (doors closed, after animal discovers closed doors), for the third criterion. We use the same place fields to determine if any new fields occurred after the door closed. We then plot spikes and behaviour for two example cells before the first spike in a new field, and after the first spike in a new field (Fig 5h), with non-local door spikes before the new field spike marked with crosses. We then regress a binary variable that indicates if a cell obtained any new place fields on a binary variable that measures if the cell replayed (i.e. had any non-local door spikes), with an additional regressor of the cell’s average firing rate (Fig 5i).

### 9. Home-well mapping simulations

To better understand the origin of the neural responses and representations that we discover in the home-away well paradigm of (Pfeiffer and Foster 2013), we simulate them through our model of compositional map making. We generate a population of synthetic conjunctive hippocampal neurons. In this population, we simulate replays that build a home well map to demonstrate the source and magnitude of replay spike aligned ratemap changes (Fig 5). We then compare simulated ratemaps before and after the home well change on day 2, to show how anticorrelated ratemap changes around the previous versus the new home well arise from compositional representations (Fig 6).

The simulated neural population consists of 10 entorhinal grid cells, 20 entorhinal object-vector cells, and 40 entorhinal cells that code for sensory observations. These entorhinal cells give rise to a population of 600 conjunctive hippocampal neurons: 10*20 = 200 landmark cells (Fig S9a,b), from the conjunction of 10 grid cells and 20 object-vector cells, and 10*40 = 400 place cells (Fig S9c), from the conjunction of 10 grid cells and 40 sensory neurons. We simulate the ratemaps for each of these neurons as follows. For each grid cell, we define grid peak locations on a triangular lattice with 120cm between peaks, rotated by 6 degrees, and add a random lattice translation per grid cell. We then calculate grid cell ratemaps by centering a 2D multivariate gaussian with a 2cm diagonal covariance matrix on each peak location. For each object-vector cell, we sample a uniform random direction and a uniform random distance between 0cm and 200cm to obtain the cell’s vector relation to the home well, then place a 2D multivariate gaussian with a 1.2cm diagonal covariance matrix at that vector from the home well. As the home well is in a different location on day 1 versus day 2 of the experiment, we generate two sets of ratemaps for the object-vector cells, one for each day. We define each sensory neuron by sampling a uniform random 2D coordinate within the arena and placing a 2D multivariate gaussian with a 2cm diagonal covariance matrix at that location. Finally, we scale the ratemaps of grid cells, object-vector cells, and sensory cells by a factor 20Hz divided by the grid/ovc/sensory maximum firing rate, so that the maximum firing rate across the population of each of those will be 20 Hz. Having generated all the required entorhinal representations, we now simulate the resulting hippocampal conjunction ratemaps simply by calculating the element-wise product of entorhinal grid cell ratemaps and object-vector cell ratemaps (for hippocampal landmark cells) and entorhinal grid cell ratemaps and sensory cell ratemaps (for hippocampal place cells). We also scale these conjunctions so they have a maximum firing rate of 20Hz across the population. Because we have two sets of ratemaps for object vector cells, due to the change in home well location on day 2, we also have two sets of ratemaps for landmark cells.

With these synthetic neural populations in place, we can now turn to simulating the ratemap changes induced by replay. Our model predicts landmark cells that appear in replay before they appear in physical navigation - in other words, their first spike happens in a replay event. To simulate that, we sample replay trajectories, and sample spikes from our synthetic hippocampal population along those trajectories. If the replay trajectory passes through the receptive field of a landmark cell for the first time, that landmark will spike, and it will have a firing field at that location for the remainder of the session. Therefore, if we calculate the change in ratemap before the replay event (no spikes) versus after the replay event (lots of spikes) for that landmark cell, we will find a large new peak (Fig S10a). The same replay trajectory may also pass through the receptive field of place cells, and these will also spike during replay, but since their representation remains stable (the sensory properties of the environment do not change), the corresponding ratemap changes will be small (Fig S10a). According to our model, we particularly expect replays from home to lay down landmark memories. We therefore contrast such replays with replays elsewhere. In this simulation, replays elsewhere may also elicit place cell and landmark cell spikes, but they will never create new landmark cells - so a landmark cell will never fire for the first time in a replay elsewhere (Fig S10b). This means that towards the end of the session, when most of the landmark cells have already been created in previous home replays, the difference between home and away replays diminishes (Fig S10c,d). Moreover, since these landmark cells have been around for a while, the difference in their ratemap before versus after the replay will be small. Therefore, when we plot the average ratemap change around replay spikes (Fig 5f), and average this change within 8cm of the replay spike (Fig 5g), we observe that 1) the map changes around both home replays and replays elsewhere (because both sample landmark spikes), 2) this change is larger for home replays (because only these sample landmark spikes for the first time, which is accompanied by the largest ratemap change), and 3) this change is much smaller than the magnitude of the new landmark firing field, because it is averages across the hippocampal population (which includes many place fields that do not change) and across replays (for which ratemap changes diminish over the course of the session).

We will now provide the implementation details of the replay simulation described above. To obtain replay spike aligned ratemap changes, we need to simulate a) replay trajectories, b) spikes of cells along these trajectories, and c) ratemap changes for those cells. We sample 100 replay trajectories, with one home replay followed by three replays elsewhere to roughly match the home/elsewhere ratio in the data. We distribute these replays evenly through time between 10s and 3000s (the length of a recording session). Each replay trajectory starts from the home location (home replays) or a random location in the arena (replays elsewhere), and extends randomly between 100cm and 300cm in a random direction, divided over 20 steps. The trajectory is cut short if it leaves the arena, or if it approaches the home well within 30cm for replays elsewhere (to match the analysis of empirical data). That gives us our replay trajectories (a). We then sample spikes for each simulated cell from a Poisson distribution where the rate is given by the simulated ratemap at the replayed locations. Since a replay step takes 5ms (as in the data), and the simulated ratemaps are modelled after those during awake behaviour, the effective firing rate in a replay event is much higher than during behaviour. To simulate this, we scale up the firing rates by a factor 25 to get around 30 replay spikes per replay trajectory, roughly matching the data. Furthermore, we assume that a landmark cell can only spike in a replay elsewhere if it has already spiked at least once in a replay from home. That gives us our replay spikes (b). The change in ratemap for a landmark cell in a replay at time t1, after it spiked for the first time at t0, is given by t0/t1 times the total ratemap change (i.e. the full new landmark peak). This implements the fact that for late replays, the landmark field already existed before that replay, so the change before versus after that replay is smaller (but for the first replay, when t1 = t0, it is the full new landmark peak). On top of that, we sample change map noise from a standard normal distribution, multiply by 0.5 Hz, and smooth this with a 4cm gaussian kernel. Place cell ratemap changes consist of only this noise. That gives us ratemap changes for each replay spike (c). With these ingredients in hand, we calculate the replay spike aligned ratemap change in the same way as we did for the empirical data.

The neural recordings by Pfeiffer and Foster did not allow for tracking the same cells over multiple days, so to find signatures of compositional vector responses, the analysis of Figure 6 relies on the total ratemap change on the second day. In our simulated neural responses, we have the luxury that we can model the same conjunctions across two days, to show that compositional conjunctive representations indeed produce the same signature as found in the empirical data. We calculate the cumulative ratemap change on day 2 for each hippocampal cell by subtracting the simulated ratemaps on the first day from the simulated ratemap on the second day, and adding change noise as in the replay simulation (Fig S9d). We then calculate the correlation between each cell’s change map aligned on the new home well and the same change map aligned on the previous home well (Fig 6d), exactly like in the empirical data analysis. We plot the distribution of correlation coefficients across the population of synthetic hippocampal neurons (Fig 6e, left), but now we know which of these are landmark cells and which are place cells, so we can separate their contributions (Fig 6e, right).

### 10. Non-spatial replay

Finally, we show how our model of state-space composition can be extended to accommodate hierarchies of spatial and non-spatial structures (Fig 7). As an example of such an environment, we consider five discrete spatial regular square grid rooms, with shapes that are randomly sampled from (4×4, 3×5, 5×3), connected on a length-5 non-spatial loop. Now transitions exist on both hierarchical levels, within-room and between-room; in general, there can be many recursive levels where a location on the higher level corresponds to a whole environment on the lower level, and a lower-level action can take the agent to a different higher-level location. Here, the low-level rooms contain two doors, which when entered transition the agent to the adjacent high-level room, and one of the low-level rooms contains reward. Crucially the doors are located randomly within each room so their location tells you nothing about the high level action. We provide the agent a state representation that consists of within-room door and reward vectors *d^1^*, *d^2^*, *r* and between-room vector *c* that specifies the clockwise and anticlockwise distance towards the rewarding room.

Like before, we train a policy mapping *f*(*s*) = *a*, where the state representation input concatenates vector representations across hierarchical levels *s* = [*d^1^*, *d^2^*, *r*, *c*] and the action output produces an ‘primitive’ action, on the lowest level of the hierarchy. Intuitively, it makes sense that this state representation affords correct policies: the *c* representation tells the agent which door to use, and the *d^1^*, *d^2^*representations tells them how to get to that door - or directly to reward following *r* if already in the rewarding room. The agent can therefore reuse the same object vector cell population in every room (Fig 7b). We implement *f* as a feedforward neural network with one hidden layer of twice the size of the input dimension. To train *f*, we generate many different environments where the shapes of the rooms, the door locations, and the rewarding room and reward location are randomised. We train *f* through supervised learning by backpropagation on 500 samples of [state representation, optimal action] pairs from 25 of such environments in parallel, and repeat this 10 times with new environments.

Since door locations are different for each room, the agent utilises a low-level memory bank for each room that binds within-room vector representations to low-level locations. Additionally, the agent has a high-level memory bank that binds between-room vector representations to high-level rooms. This setup factorises the different levels of the hierarchy, and so replay can be independent for the different levels of the hierarchy.

When the agent has, through latent learning, built low-level memories of all door-vectors *d^1^*, *d^2^*within every room before finding a reward, it has complete maps of the low-level environments but no optimal policy yet - it knows where the doors are, but not yet which door to take. But when it finds the reward in the rewarding room, it only needs to replay at the high-level to create a memory that binds the between-room reward vector *c* to the room to get access to the optimal policy (Fig 7c). Next time when it enters a room, it retrieves *c* from high-level memory and *d^1^*, *d^2^*from the low-level memory bank for that room, which tells it both where the doors are and which door to approach.

### 11. Summary in double dactyl

*Lego with state-spaces: Cortical building blocks bound through conjunction in any new way*.

*Thus hippocampus is combinatorially building behaviour like children’s (re)play*.

## Footnotes

1 We refer to the cortical building blocks as *compositional* as they can occur in any configuration, but we propose the *composition* of these building blocks occurs in hippocampus.

## Notes

### Competing Interest Statement

The authors have declared no competing interest.

### Summary of Updates

New section on empirical validation of our proposed model in hippocampal recordings

