## Supplementary Material for "Constructing future behaviour in the hippocampal formation through composition and replay"

November 8, 2023

### 1 Model implementation details

#### 1.1 Parameters

| Name | Value | Description |
|---|---|---|
| $l_d$ | 7 | Number of nodes along one side of square grid graph |
| $n_b$ | 25 | Number of training samples in batch, each from a different environment |
| $n_s$ | 200 | Number of training samples from the same environment |
| $n_e$ | 20 | Number of times new training environments are generated |
| $\epsilon$ | 0 | Path integration error probability |
| $w$ | 0.2 | Weight of path integrated representation (versus memory retrieved representation) in representation update |
| $r_n$ | 5 | Number of replays repeats at one step |
| $r_l$ | 15 | Replay length: number of steps in one replay |
| $r_f$ | 4 | Replay interval: number of physical steps after which to replay |
| $l_c$ | 1 | Length of continuous two-dimensional arena side |
| $n_c$ | 100 | Number of object vector cells in object vector representation of one object |
| $d_R$ | 0.1 | Maximum step size |
| $\sigma_r$ | 0.01 | Step size variation standard deviation |
| $\sigma_{theta}$ | $0.1{\cdot}2\pi$ | Step direction variation standard deviation |
| $l_w$ | 0.025 | Wall radius (or half the wall thickness) |
| $r_o$ | 0.05 | Radius for perceiving objects |
| $\epsilon_r$ | 0.025 | Path integration step size noise standard deviation |
| $\epsilon_\theta$ | $0.25{\cdot}2\pi$ | Path integration step direction noise standard deviation |
| $\xi_M$ | 0.05 | Distance cutoff for including memories in retrieval |
| $\beta_M$ | 1 | Memory retrieval weight softmax inverse temperature |

List of parameters with default values. These values are not fixed across simulations: often a simulation systematically varies one or more of these parameters (e.g. the path integration noise, or the number of replays).

#### 1.2 State representation

##### 1.2.1 Discrete model

The state space of the discrete model is defined on a set of states $s \in \mathcal{S}$, connected by actions $a \in \mathcal{A}$, to form a graph as in a deterministic Markov decision process. All spatial environments are regular square grid graphs, with actions 'go-north', 'go-east', 'go-south', and 'go-west'. We generate environments by populating a $l_d \times l_d$ square grid with walls and rewards: we assign nodes a 'wall' or 'reward' identity. Wall nodes can be horizontal or vertical neighbours and cannot overlap with rewards or completely block off parts of the environment. We disconnect wall nodes from their non-wall neighbours in the environment by disabling actions that would transition the agent to the wall node.

The full compositional state representation $\vec{v}$ of state $s$ consists of the concatenation of the vector representations $\vec{v}_k$ across all components $k$ in the environment. We represent a reward as a single component, and a wall as a pair of components, one on each end of the wall. For example, the full representation of a state in an environment with two walls and one reward will be $\vec{v} = [\vec{v}_{w_1 1}, \vec{v}_{w_1 2}, \vec{v}_{w_2 1}, \vec{v}_{w_2 2}, \vec{v}_r]$ where [] means concatenation and $\vec{v}_k$ is a vector representation. We implement this vector representation $\vec{v}_k$ as follows (Figure 1). For state $s$, we get the representation $\vec{v}_k$ of component $k$ at state $q$ as the concatenation of one-hot encoded distances along all actions from $s$ to $q$ (Figure 1a). We set the distance to -1 for an action if $q$ is in the opposite direction of that action (Figure 1b). For example, if $q$ is 1 step to the east and three steps to the south of $s$, the state representation $\vec{v}_k$ at $s$ becomes $[onehot(-1), onehot(1), onehot(3), onehot(-1)]$ (Figure 1c,d). Here [] means concatenation and $onehot(i)$ is the (i+2)-th column of the $(l_d + 1)$-dimensional identity matrix: each one-hot vector has dimension $l_d + 1$, to enable encoding of all possible distances in the square grid environment, plus the -1 for opposite direction in the first dimension. The resulting $(4\ l_d + 4)$-dimensional vector thus always contains 4 ones, one for each action. Finally, in addition to the vector representations for all components in the environment, we implement a one-hot place code (our model's analogy to the brain's grid code - a coordinate system that uniquely identifies each state) for state $s$ as the $i_s$-th column of the $(l_d \cdot l_d)$-dimensional identity matrix.

##### 1.2.2 Continuous model

Continuous environments are specified on two-dimensional continuous $(x, y)$ coordinates in an $l_c \times l_c$ arena, so $\vec{s} = \{(x, y)|0 \leq x \leq l_d; 0 \leq y \leq l_d\}$. We place reward and walls within this arena (Figure 2). Again, a reward is represented by a single vector code, and a wall by two vector codes, one centred on each wall end (Figure 2a,b). And like the discrete environment, the full compositional representation for a state concatenates the vector representations for all components, e.g. $\vec{v} = [\vec{v}_{w_1 1}, \vec{v}_{w_1 2}, \vec{v}_{w_2 1}, \vec{v}_{w_2 2}, \vec{v}_r]$. But the individual vector representation $\vec{v}_k$ now consists of the firing rates of a population of $n_c$ spatially tuned object-vector cells (Figure 2c,d). For a component $k$ at state $\vec{q}$, we distribute the firing

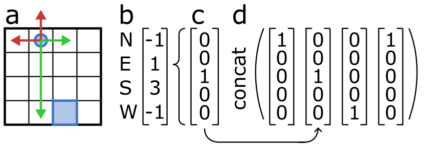


Figure 1: **Discrete object vector representations.** a. Example object vector representation at state $s$ indicated with the blue circle for the blue square reward component at state $q$. We calculate distances along each action to the component, plotted by green arrows; red arrows indicate actions that go in opposite direction. b. The full object vector representation is a vector of distances along each action, here 'go-north' (N), 'go-east' (E), 'go-south' (S), and 'go-west' (W). Opposite actions are indicated by a -1. c. We one-hot encode each action distance. d. The full state representation $\vec{v}$ that serves as input for $f$ is the concatenation of these one-hot encoded distances.

fields of the object-vector cells evenly on a $2 \cdot l_c \times 2 \cdot l_c$ square grid, with the population's centre aligned on $\vec{q}$. This configuration makes sure that the whole arena is covered by object-vector cells for any possible $\vec{q}$. We model the firing field of a single object-vector cell as a two-dimensional multivariate Gaussian, with a diagonal covariance matrix that has firing field width $\Sigma$ on the diagonal and zeros off-diagonal. Then to obtain the vector representation $\vec{v}$ at state $\vec{s}$ for element $k$, we evaluate the $\vec{q}$-centred population of $n_c$ multivariate Gaussians at $\vec{s}$ to get a $n_c$-dimensional vector of activities.

Like in the discrete environment, we also implement a place code that represents the absolute location of a state, analogous to grid cells that encode a coordinate system in 2d space to uniquely index location (Figure 2e). In the continuous environment, this place code is obtained by evaluating $n_c/4$ multivariate Gaussians that are arranged at fixed locations on a regular square grid within the arena.

##### 1.2.3 Conjunction versus concatenation

Throughout this paper, state representations are sometimes written as conjunctions and sometimes as concatenations. To avoid confusion, we'll distinguish the two as explicitly as possible here. A concatenation creates a new representation that stacks its component representations; the dimension of the new representation will be the sum of its component dimensions. We use concatenated representations to produce policies from a set of vector representations (e.g. a reward-vector code and a wall-vector code): we map the concatenated vector representations to actions (Section 1.3). A conjunction creates a new representation that combines its component representations, for example through an outer product; the dimension of the new representation will be the product of its component dimensions. We use conjunctions to bind vector representations (e.g. a reward-vector code) to place representations (e.g. a grid code) and store this combination in memory (Section 1.4). Importantly, these conjunctive

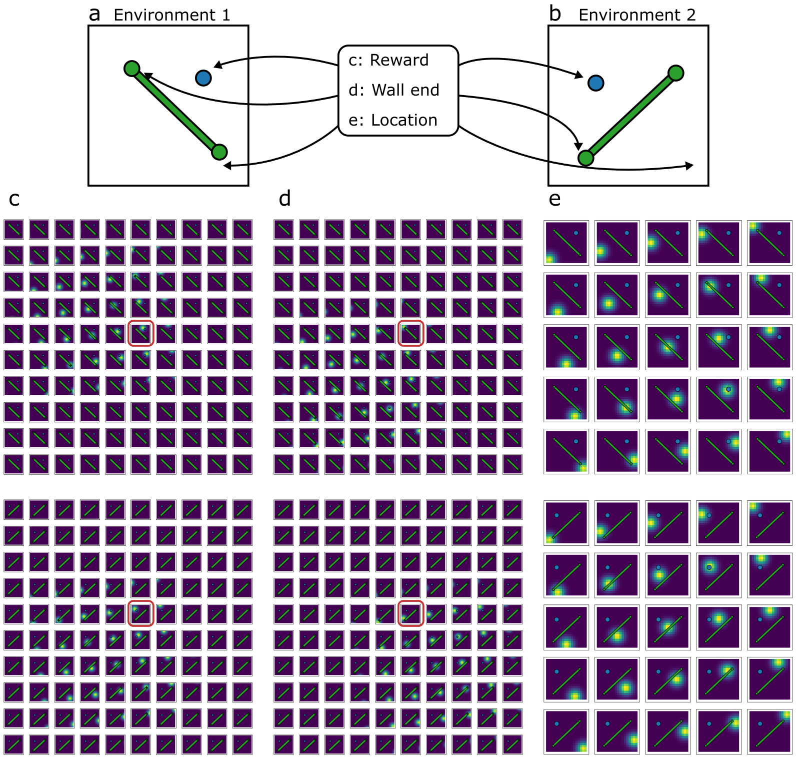


Figure 2: **Continuous object vector representations.** a. Continuous environment with a reward (blue circle) and a wall (green bar), represented by two wall ends (green circles). b. Continuous environment with different configuration. c. Object vector cell population for the reward object. Each square is a rate map for one cell, the top row for the environment in *a* and the bottom row for the environment in *b*. The rate map circled in red, for example, shows a cell that fires just west of the reward in both environments. d. Object vector cell rate maps for one wall end, with the cell circled in red firing just west of the wall end in both environments. e. Place cell population that represents location in both environments, independent of object orientation.

memories can be created independently for each component: the agent can create a map of grid-reward conjunctions, and a map of grid-wall conjunctions, at different times and different places. Then when they need to select an action at a particular location, encoded by the grid code, they can retrieve the corresponding vector codes, e.g. the reward-vector and wall-vector, concatenate them, and infer appropriate action (Section 1.5). In summary: in our model conjunctions are for memory, and concatenations are for policy.

#### 1.3 Learning policies that generalise

##### 1.3.1 Discrete model

Given the full state representation that concatenates vector codes for all objects in the environment, e.g. $\vec{v} = [\vec{v}_{w_1 1}, \vec{v}_{w_1 2}, \vec{v}_{w_2 1}, \vec{v}_{w_2 2}, \vec{v}_r]$, the model learns a policy through a mapping $f(\vec{v}) = \vec{p}_a$ that maps a state representation to action probabilities. Here $\vec{p}_a$ is a vector of action probabilities, so its dimension matches the number of actions in the environment, i.e. four in the square grid world. The mapping $f$ is learned across many environments, so that it generalises to new configurations of objects on first encounter. We implement $f$ as a multi-layer perceptron, consisting of an input layer of the same dimension as $s$ (i.e. $(4\ l_d + 4)$*number of components), then three hidden layers of dimensions 1000, 750, 500 and rectified linear activation, and an output layer of the same dimension as $a$ (i.e. 4). Before inputting $\vec{v}$ into $f$, we preprocess all wall vector representations with a common wall-embedding function $f_w$, with an output layer of the same dimension as the wall representation and a hidden layer twice that size and rectified linear activation. In the example $\vec{v} = [\vec{v}_{w_1 1}, \vec{v}_{w_1 2}, \vec{v}_{w_2 1}, \vec{v}_{w_2 2}, \vec{v}_r]$, that means that the input to $f$ would be $[f_w(\vec{v}_{w_1 1}, \vec{v}_{w_1 2}), f_w(\vec{v}_{w_2 1}, \vec{v}_{w_2 2}), \vec{v}_r]$. This wall embedding is not necessary for accurate policies, but does speed up learning.

We train $f$ and $f_w$ through supervised learning on the optimal policy. As training examples, we generate $n_b$ environments with random configurations of walls and rewards, and calculate the representation $s$ and optimal policy $a$ for each state. We use Dijkstra's algorithm to determine shortest-path distances from each location to the reward, while taking walls into account; the optimal policy then assigns equal probability to all actions that decrease the distance to reward, and zero to others. We then sample a random node from each environment and train $f$ on the batched $\{\vec{v}, \vec{p}_a\}$ pairs. We update $f$'s parameters through backpropagation using gradients provided by Pytorch and the ADAM optimiser with learning rate $10^{-3}$. Because $f$ is trained on multiple environments in parallel, it needs to learn a mapping that works across environment configurations. After $n_s$ samples we generate new training environments, and this procedure is repeated $n_e$ times.

##### 1.3.2 Continuous model

In continuous environments, we define actions as optimal directions to reward. Therefore mapping $f(\vec{v}) = \vec{\phi}$ must produce a continuous, periodic value $-\pi \leq \theta \leq \pi$. We implement this action output as a two-dimensional vector $\vec{\phi}$ that contains the sine and the cosine of the optimal direction $\theta$; the optimal direction can be recovered from this output as $\theta = atan2(\phi_2, \phi_1)$. Again, the input state $\vec{v}$ concatenates the vector representation across all components in the environment, e.g. $\vec{v} = [\vec{v}_{w_1 1}, \vec{v}_{w_1 2}, \vec{v}_{w_2 1}, \vec{v}_{w_2 2}, \vec{v}_r]$ – with vector representations calculated as described in Section 1.2.2. The network hidden state dimensions are 3000, 2000, 1000, and we preprocess wall embeddings as in the discrete model to speed up learning. As training examples, we generate $n_b$ environments with random configurations of walls and rewards, and sample random locations within those to provide training state-action pairs. We calculate the optimal direction $\theta$ for a location as follows. First, we construct a fully connected graph of the environment, where each component in the environment (with two components for a wall, one on each end) is a node, and with edges whose weight is the Euclidian distances between objects. We remove edges between objects if the line connecting them in the environment crosses any walls. Then we use Dijkstra's algorithm to calculate the graph distance to the nearest reward for each node on this graph, taking the edge weights into account. Finally, for a given location $\vec{s}$, the optimal direction points towards the object that a) can be reached in a straight line without crossing walls from $\vec{s}$ and b) has the smallest graph distance to reward after adding the distance from that object to $\vec{s}$. Provided with these state-action training examples, we update $f$'s parameters through back-propagation as in the discrete model. After $n_s$ state-action training examples we sample new training environments, and we repeat this $n_e$ times.

#### 1.4 Mapping the environment

##### 1.4.1 Discrete model

Given a learned mapping $f(\vec{v}) = \vec{p}_a$, the agent can act optimally when it has access to its full state representation $\vec{v}$. However, upon entering a new environment, they don't know the current configuration of components. They will need to incorporate the components in their state representation through exploration (Figure 3). A simulated episode in such a new environment runs as follows. Initially, all vector representations are initialised as empty. The agent explores the environment by observing their location, selecting an action from their exploration policy (e.g. a random policy with equal probability for all available actions), and transitioning to the next location. When they discover a component, by approaching it within one step for wall ends or zero steps for reward, they initialise the object-vector representation $\vec{v}_k$ for that object $k$. For ease of notation and implementation, we will keep track of the represented state $s_k$ of the agent for that component rather than the vector representation $\vec{v}_k$. This representation can be thought of as "where does the agent think they are, with respect to component $k$". For example, when the agent discovers a component $k$ at state $q = 12$, and takes a step east, we'll write $s_{k,q=12} = 13$ for their new vector representation of that component instead of $\vec{v}_k = [onehot(0), onehot(-1), onehot(0), onehot(-1)]$ ('1 step west'). That makes the equations easier to read, but the two notations are equivalent, because there is a one-to-one correspondence between represented state and object-vectors (as described in Section 1.2.1). We'll drop the subscript $q$ in the following because for a particular component $k$, the state it is in, $q$, is fixed.

After initialisation, the agent carries out such updates to vector representations with respect to their actions through path integration (Figure 3a). However, path integration is noisy. Errors may cause the represented component vector state $s_k$ to diverge from the true state $s$. We model path integration noise as a distribution over transitioned locations $s'_k$ given action $a$ from state

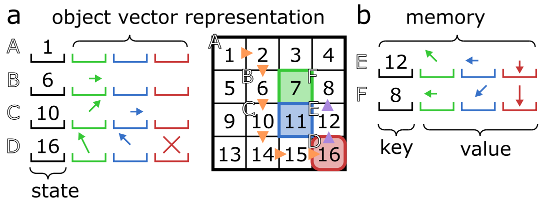


Figure 3: **Discrete map-making.** a. Schematic of a model trajectory after entering a new environment for the first time. During exploration (orange arrows), the model observes states (numbers, black slots) and tracks object vector representations (arrows in green, blue, and red slots) through path integration. Vector representations start empty (step A), but are initialised on component discovery (like the green wall end, step B). As the agent moves from state 6 to state 10, they uses their knowledge about their actions to path integrate the representations of the green wall (so that now in state 10 it points north-east). In addition, the model discovers the blue wall, and initialises its representation. b. At each step, the model stores the vector representations it has access to in memory by binding it to the state, implemented as a dictionary where the state (black) is the key and the representation (green, blue, red) the value. In replay, the model carries state representation to remote locations without physical displacement (purple arrows). It then binds the object vector representations to locations in memory (step E, F) for later retrieval at that location.

$s_k$:

$$p_{PI}(s'_k|s_k, a) = \begin{cases} \frac{1-\epsilon}{|S'(s_k,a)|} & \text{if } s'_k \in S'(s_k, a) \\ \frac{\epsilon}{|S''(S'(s_k,a))|} & \text{if } s'_k \in S''(S'(s_k, a)) \\ 0 & \text{otherwise} \end{cases} \tag{1}$$

where

$$S'(s_k, a) = \{s' \in \mathcal{S} \mid p(s'|s_k, a) > 0\} \tag{2}$$

is the set of locations that taking $a$ from $s_k$ can transition to ($p(s'|s, a)$ is defined by the environment; here, since the environment is deterministic, $|s'| = 1$), and

$$S''(S') = \{s'' \in \mathcal{S} \mid s'' \notin S' \text{ and } \sum_{s'}^{S'} \sum_{a'}^{\mathcal{A}} p(s''|s', a') > 0\} \tag{3}$$

is the set of two-step neighbours of $s_k$. In other words, path integration updates the representation to the correct location $s'_k$ with probability $1 - \epsilon$ or any of the correct location's neighbours with total probability $\epsilon$. Note that this accumulation of error occurs independently for each component $k$, resulting in a (possibly) different representation of "where do I think I am" with respect to each component.

In addition to path integration, the agent relies on memory to update representations after an action (Figure 3b). In the brain, we hypothesise these memories are stored as hippocampal conjunctions, which link a vector representation (the object-vector cell population activity) to place representations (the grid cell population activity).

Here, we model such conjunctive memories as a dictionary $M_s = [s_k^i, ...]$ with states $s$ as keys and multisets (i.e. the same element can appear multiple times) of vector representations (or, again for ease of notation: the state $s_k$ corresponding to a vector representation) as values. The superscript $i$ indexes all the memories that exist at state $s$: the agent appends a new memory $s_k$ to the memory bank $M_s$ at state $s$ every time they go there. Without path integration errors, the represented state $s_k$ always matches the true state $s$, so in that case $M_s$ would just contain many copies of $s$. But due to path integration errors, the represented state might diverge from the true state, so that the list of memories $M_s$ contains a variety of states $s_k$ that are not $s$. In the example of taking a step east after finding reward at state 12, the new memory would be $M_{13} = [13]$, indicating a reward one step to the west at state 13 – if they have correctly path integrated their reward-vector representation. If they made a path integration error, the encoded memory may instead be $M_{13} = [14]$, (incorrectly) indicating a reward two steps to the west. When the agent returns to a location where it has previously encoded memories, they can retrieve these memories. We model the probability of retrieving a particular vector representation $s_k$ as proportional to the number of encoded memories of that vector representation at the current state $s$:

$$p_M(s_k|s) = \frac{|[s_M \in M_s \mid s_M = s_k]|}{|M_s|} \tag{4}$$

which is only calculated if $|M_s| > 0$, so there are memories at state $s$. Equation 1 and 4 thus provide two probability distribution of updated vector representations $s'_k$ given the previous representation $s_k$, the action the agent took $a$, and the new state $s$. To update their representation, the agent can make use of the combination of the two: a weighted sum over the path-integrated representation (weight $w$) and the distribution of representations on earlier encounters of this state (weight $1 - w$), setting $w = 1$ if $|M_s| = 0$.

$$p(s'_k|s_k, a, s) = w p_{PI}(s'_k|s_k, a) + (1 - w) p_M(s'_k|s) \tag{5}$$

They sample $s'_k$ from Equation 5 to obtain the updated representation, which is encoded in memory and path-integrated in the next step.

Replay in our model allows the agent to encode remote memories offline. Given an initial state $s$ and representation $s_k$, they imagine transitions through the environment. After each transition, they update their representations and append them to the current state's memories, as during online navigation. There is one difference: Since transitions are now imagined, the current state cannot be observed from the environment. Instead, it needs to be path integrated, just like the object-vector representations. That means there can now be errors both in the state $s$ (the key in the memory dictionary) and the vector representation (the value in the memory dictionary). Again, we model path integration noise by sampling from a distribution over correct and neighbour locations. The agent performs $r_n$ replays of $r_l$ steps long, every $r_f$ steps of exploration.

##### 1.4.2 Continuous model

During simulated exploration, the continuous space agent takes steps d of length $d_r$ (capped at $d_R$) in direction $d_\theta$ by adding normal random variation with standard deviation $\sigma_r$, $\sigma_\theta$ to the previous length and direction for smooth trajectories. Compared to the discrete environments, where we simply disabled actions that lead into walls, the interaction with continuous walls is a bit more complicated. The agent cannot pass through walls, but should still be able to slide along walls if they are not moving perpendicularly towards the wall, unless there is another wall preventing that (as in corners). Algorithm 1 resolves an attempted step $\vec{d}$ from state $\vec{s}$ to $\vec{s}'$ in the presence of walls, with wall normal vectors pointing from $\vec{s}$ orthogonally into the wall.

**Algorithm 1** Transition with walls

**1.** Is $\vec{s}$ within distance $l_w$ of a wall?

- **Yes** Is $\vec{s}$ within distance $l_w$ of multiple walls?
  - **Yes** Calculate the dot products of $\vec{d}$ and the normal vectors of the walls within $l_w$
    Remove the positive component along the normal vector of the wall with the largest dot product from $\vec{d}$.
    Recalculate the dot product of $\vec{d}$ and the normal vector of the wall that had the second-largest dot product. Is it greater than zero?
    - **Yes** Set $\vec{d}$ to $\vec{0}$; *continue to 2.*
    - **No** *continue to 2.*
  - **No** Remove the positive component along the normal vector of the wall within $l_w$ from $\vec{d}$; *continue to 2.*
- **No** *continue to 2.*

**2.** Does the line from $\vec{s}$ to $\vec{s} + \vec{d}$ cross any walls?

- **Yes** Set new location $\vec{s}'$ to wall contact location, $l_w$ away from the wall crossed first; *exit.*
- **No** Set new location $\vec{s}'$ to $\vec{s}' = \vec{s} + \vec{d}$; *exit.*

The agent thus explores the environment (Figure 4). They discover a component when they get within radius $r_o$ of that component, and initialise the corresponding vector representation upon discovery. Again, for ease of notation, we will write the agent's represented state $\vec{s}_k$ for an object instead of the object-vector representations $\vec{v}_k$. In other words, we'll write representations as 2-dimensional $(x, y)$ vectors, but these have a one-to-one correspondence with $n_c$-dimensional population activities of object-vector cells. After initialisation, the agent tracks the object's representation by combining path integration with

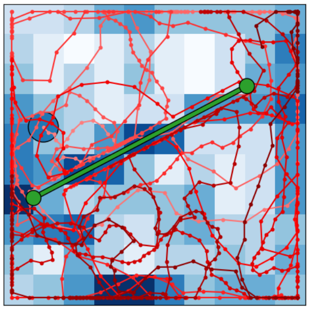

Figure 4: **Continuous exploration.** Example walk of agent in a continuous environment, each step marked with a red dot connected to the previous step with a red line, progressing from light red to dark red over the course of the walk. Blue patches indicate binned location occupancy. Interactions with the green wall and arena boundaries demonstrate how the agent slides along blockades without crossing them.

memory retrieval, just like in the discrete model. Path integration updates represented states $\vec{s}_k$ to reflect step $\vec{d}$, with path integration error modelled as Gaussian noise with standard deviation $e_r$ and $e_d$ on step length $d_r$ and direction $d_\theta$ respectively:

$$
\begin{aligned}
\vec{s}_k^{PI} &= \vec{s}_k + \vec{d}^{PI} \\
\vec{d}^{PI} &= d_r^{PI} \cos(d_\theta^{PI})\hat{x} + d_r^{PI} \sin(d_\theta^{PI})\hat{y} \\
d_r^{PI} &= d_r + e_r, e_r \sim \mathcal{N}(0, \epsilon_r) \\
d_\theta^{PI} &= d_\theta + e_\theta, e_\theta \sim \mathcal{N}(0, \epsilon_\theta)
\end{aligned}
\tag{6}
$$

Note that equation 6 follows the state update algorithm, taking walls into account.

We encode a new memory as a state-representation pair, so that the memory bank $M = [\{\vec{s}^i, \vec{s}_k^i\}, ...]$ becomes a multiset of pairs. The reason this is different from the dictionary in the discrete case, is that states are now continuous coordinates, so there is not much point in using them as dictionary keys: it is unlikely the same key will occur twice. Again, this memory bank is a simplified implementation of what we hypothesise the brain achieves through hippocampal conjunction: the binding together of a location and object-vector code. Then to retrieve a representation at state $\vec{s}$ from all previous experience, we calculate a weighted average across representations $\vec{s_k}^i$ in the memory bank. The weights are given by the corresponding memory states $\vec{s}^i$: memories created near $\vec{s}$ contribute stronger to the retrieved representation. More specifically, we exclude memories create beyond distance threshold $\xi_M$, and then weight remaining memories by the softmax of dot products between the memory locations $\vec{s}^i$

and retrieval location $\vec{s}$:

$$\vec{s}_k^M = \frac{\sum_{\{\vec{s}^i, \vec{s}_k^i\} \in V} e^{\beta_M \vec{p}(\vec{s}) \cdot \vec{p}(\vec{s}_i)} \vec{s}_k^i}{\sum_{\{\vec{s}^i, \vec{s}_k^i\} \in V} e^{\beta_M \vec{p}(\vec{s}) \cdot \vec{p}(\vec{s}_i)}} \tag{7}$$

where $\beta_M$ is the softmax inverse temperature, $\vec{p}(\vec{s})$ the place code of a given $(x, y)$ location (see Section 1.2.2) and $V$ only contains the included memories:

$$V = [\{\vec{s}^i, \vec{s}_k^i\} \in M \mid |\vec{s} - \vec{s}^i| < \xi_M] \tag{8}$$

The final updated representation combines the path-integrated and retrieved representation as a weighted sum, again setting $w = 1$ if $|V| = 0$:

$$\vec{s}_k = w\vec{s}_k^{PI} + (1 - w)\vec{s}_k^M \tag{9}$$

This final updated representation is encoded in memory and path-integrated in the next step. The replay implementation in the continuous model is very similar to the discrete case. The agent imagines transitions and makes a new memory at each replay step, path-integration both location and representation – in contrast to physical navigation, where the location $\vec{s}$ is observed from the environment.

##### 1.5 Goal-directed navigation

Following the map construction of Section 1.4, the agent can read out the policy given a state representation through the policy mapping learned as described in Section 1.3. By carrying out these actions, the agent engages in goal-directed navigation or exploitation, in contrast to Section 1.4 which described exploration. Practically, we first convert the represented locations to vector codes for each object as described in Section 1.2, and then concatenate the vector codes across objects to obtain the input vector for $f$. The agent then executes the action (in the continuous model, this means taking a step of size $d_R$ in the direction produced by $f$), updates their representations, and repeats the procedure until reaching reward.

#### 2 Supplementary figures

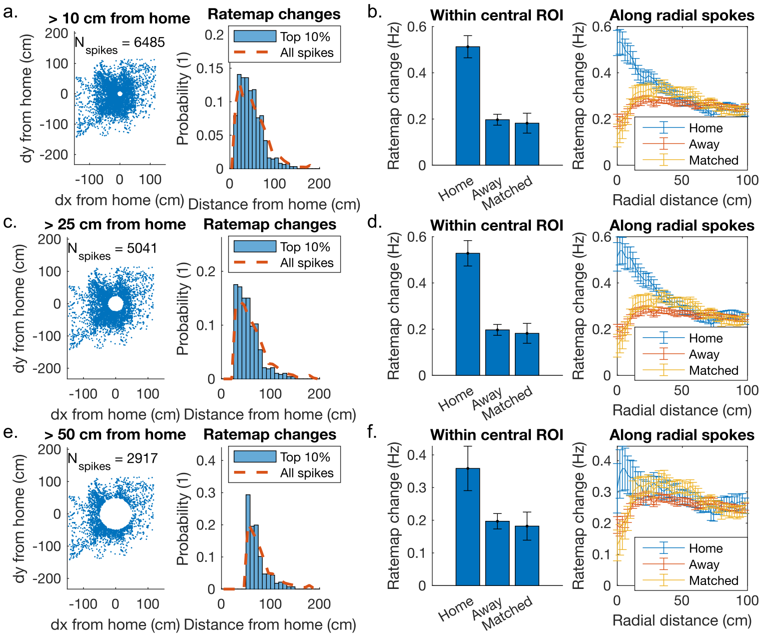


Figure 5: **Contribution of remote replay spikes.** a. To verify that the replay-spike aligned ratemap changes are not just driven by changes at the home-well, we repeat the analysis while excluding replay spikes within 10 cm from the home well. The location of included replay spikes relative to home is plotted on the left, and the distribution of home distances for the top 10 % of replay spikes that change the ratemap the most on the right, showing that many ratemap changes occur at considerable distance from the home well. The baseline distribution of home distances across all replay spikes is shown on top in dashed lines. b. Repeating region-of-interest and radial-spoke averages of ratemap change around replay spikes, only including replay spikes more than 10 cm away from the home well as shown in *a*. c-d. Same procedure, now excluding replay spikes within 25 cm from the home well. e-f. Same procedure, now excluding replay spikes within 50 cm from the home well. As the full arena is 200 by 200 cm, the diameter of the exclusion region now spans half of the arena length.

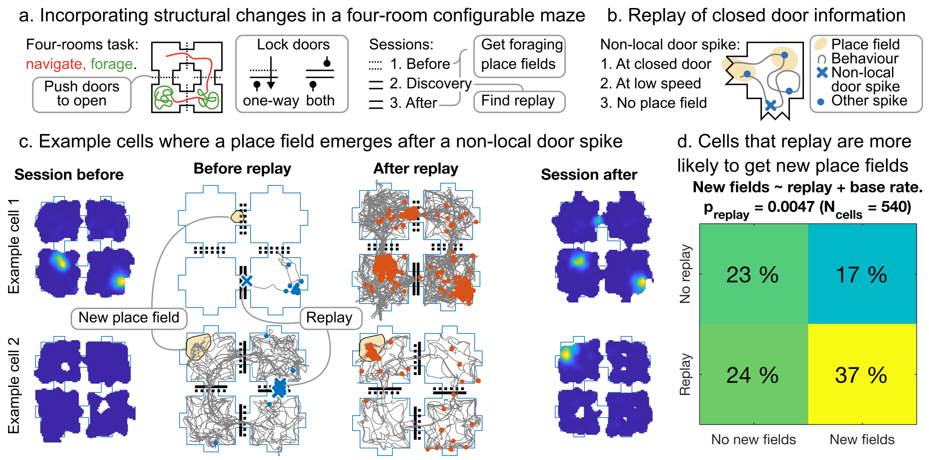


Figure 6: **Replay to incorporate structural elements.** a. We investigate replay of structural changes in a four-room configurable maze task, where the animal needs to navigate to a particular room then forage there. Some doors between rooms close halfway through the experiment; we calculate ratemaps before and after closing and look for replay after discovery of the change. b. Because we cannot detect SWRs or decode replay, we define replay through 'non-local door spikes'. c. Two example cells where such a non-local door spike precedes the emergence of a new place field. d. Across the population, cells with non-local door spikes are more likely to get new place fields. Each recording consists of five sessions: doors open (s1), doors open (s2), doors closed (s3), doors closed (s4), doors open (s5). We find cells that have non-local door spikes when the doors close for the first time (s3) and ask whether the same cells obtain new place fields in the next closed door session (s4) that did not exist in the previous open door session (s2). We regress the emergence of new fields on the presence non-local door spikes, including a regressor for baseline firing rate, and find a significant relation.

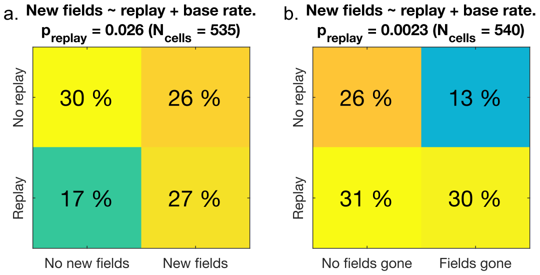


Figure 7: **Replay for stable environments and erasing fields.** a. The relation between replay and new fields also exists in a stable environment, but there is less replay than in a changing environment. We repeat the analysis of Figure 6, but now extract non-local door spikes in session s2 (a stable session), and detect new place field by comparing session s3 to session s1. The relation between the two still holds, but this time there are more cells that have no replay and no new fields (top left) and fewer that have replay and new place fields (bottom right). That is consistent with the model: even in the absence of structural changes, when replay occurs it should still make new landmark cells (to further consolidate the representation). It should just do so less often. This means that the correlation between replay and new cells should still exist, but there should be less replay than in 6, where the environmental structure changes. b. There is also a relation between replay and disappearing fields in a changing environment, but that leaves more replays unexplained. We repeat the same analysis of Figure 6, but focus on disappearing place fields rather then new place fields. We extract non-local door spikes in session s3, but detect place fields that are present in session s2 but not any more in session s4. We find that cells that replay in s3 lose place fields more often, which is consistent with our findings of changes in representations on the second day of the home-away well paradigm. This time the proportion of cells that does replay but does not change firing fields fields (bottom left) is larger than in *a* and 6, which again aligns with our model: we expect many of these replaying cells to gain new place fields too (rather than just lose them), which this analysis does not measure.

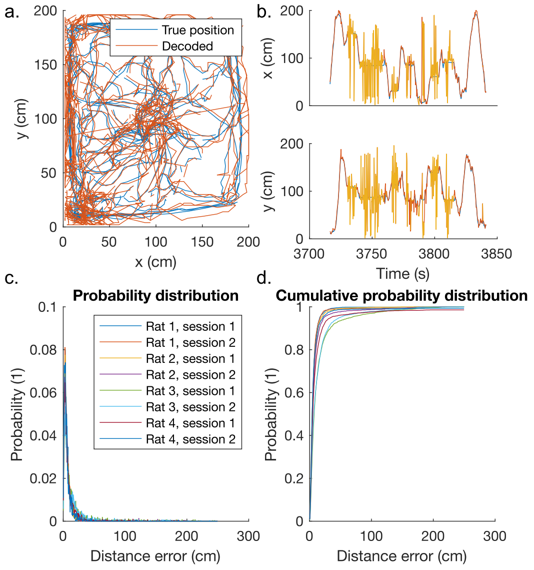


Figure 8: **Decoding accuracy.** a. Decoded trajectory from held-out neural data, plotted on top of the true position of the animal, for an example rat in an example session. This only includes time windows where animal moved faster than 5 cm/s b. Same data as in *a*, but only the first 125 seconds, separated for the x and y position, plotted against time. During periods of immobility (animal speed below 5 cm/s; flat line in true position against time), the decoded position is overlaid in yellow. c. Probability density for decoding error, measured as distance between true and decoded position, across animals and sessions. d. Cumulative probability distribution for data in *c*.

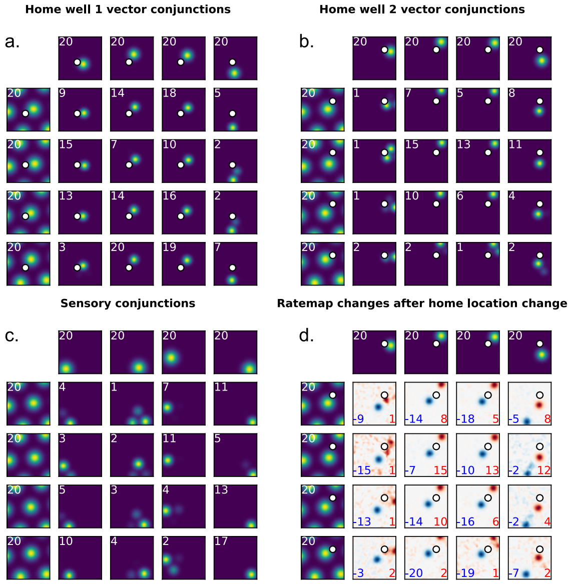


Figure 9: **Simulated ratemaps for home-well experiment.** a. Sample of synthetic entorhinal grid/object-vector cells, and hippocampal landmark cells in the simulation of the home-away well paradigm for the home well location (white circle) of day 1. The first row shows an example subset of 4 object-vector cells (there are 20 in the simulated population), the first column shows an example subset of 4 grid cells (there are 10 in the simulated population), and the remaining rows and columns the resulting 16 conjunctions (landmark cells; there are 200 of those in the simulated population). The white number indicates the max firing rate in Hz. b. Same cells as in *a* but for the home well location (white circle) of day 2. c. Sample of synthetic enthorinal grid/sensory cells, and hippocampal place cells in the same simulation. Like *a* and *b*, except that the top row is now a subset of 4 sensory cells (there are 40 of those in the simulated population), so that the 16 example hippocampal conjunctions (there are 400 in the full simulated population) are now place cells that are stable across days, because the sensory properties of the environment remain identical. d. Ratemap changes for the example subset of landmark cells in *a* and *b* from day 1 to day 2. Positive ratemap changes are plotted in red, negative ratemap changes in blue, with the red and the blue number in the right and the left bottom corner indicating maximum and minimum change respectively.

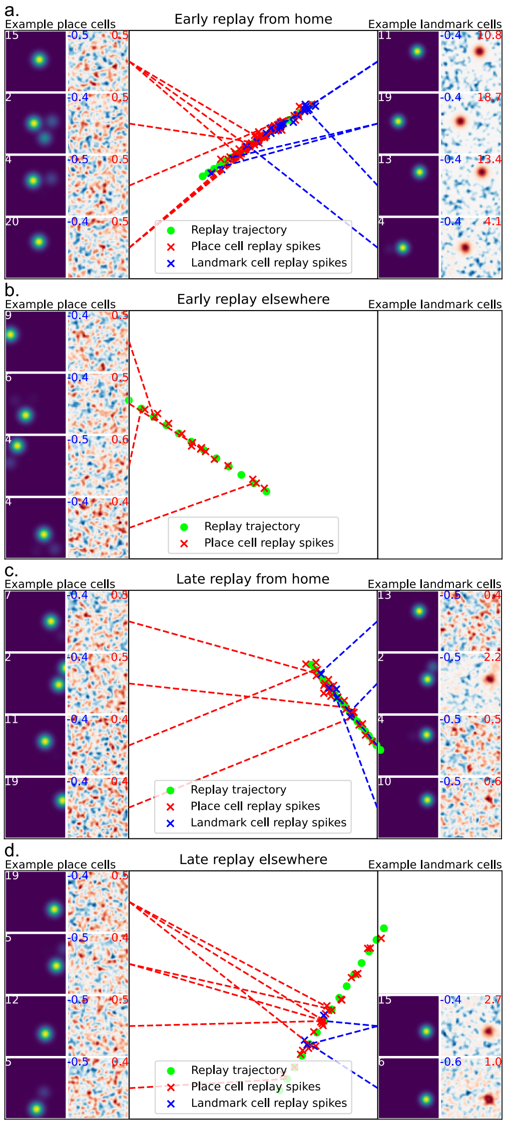


Figure 10: **Example replays in home-well simulation.** a. Example of a simulated replay from home early in the session. The green trajectory plots the replayed trajectory. Red crosses are spikes of place cells during this replay (with small jitter for illustration; else they would all be on top of the green circles, as these are the replayed locations), blue crosses are spikes of landmark cells. (Continued on next page.)

Figure 10: (Continued from previous page.) On the left of the arena there is a random sample of four place cells (left: ratemap, peak firing rate (Hz) in white; right: change in ratemap, min/max change in firing rate (Hz) in blue/red) that fired during this replay; the dashed red lines connect these cells with their replay spikes. There are four random landmark cells that fired on the right of the arena, connected to their spikes with blue dashed lines. Notice large ratemap changes for the landmark cells as these firing fields have just created in this replay. b. Example of simulated early replay elsewhere. Notice that no landmark cells fired in this replay, as their fields have not been created yet. c. Example of simulated late replay from home. Notice the small changes for landmark cells, because their fields have existed for a while, so the ratemap before this replay ends up highly similar to the ratemap after this replay. d. Example of simulated late replay elsewhere. Notice that, in contrast to *b*, there are landmark cell spikes, but again the landmark ratemap changes are small.
